# Insulin dissociates by diverse mechanisms of coupled unfolding and unbinding

**DOI:** 10.1101/2020.03.16.993931

**Authors:** Adam Antoszewski, Chi-Jui Feng, Bodhi P. Vani, Erik H. Thiede, Lu Hong, Jonathan Weare, Andrei Tokmakoff, Aaron R. Dinner

## Abstract

The protein hormone insulin exists in various oligomeric forms, and a key step in binding its cellular receptor is dissociation of the dimer. This dissociation process and its corresponding association process have come to serve as a paradigms of coupled (un)folding and (un)binding more generally. Despite its fundamental and practical importance, the mechanism of insulin dimer dissociation remains poorly understood. Here, we use molecular dynamics simulations, leveraging recent developments in umbrella sampling, to characterize the energetic and structural features of dissociation in unprecedented detail. We find that the dissociation is inherently multipathway with limiting behaviors corresponding to conformational selection and induced fit, the two prototypical mechanisms of coupled folding and binding. Along one limiting path, the dissociation leads to detachment of the C-terminal segment of the insulin B chain from the protein core, a feature believed to be essential for receptor binding. We simulate IR spectroscopy experiments to aid in interpreting current experiments and identify sites where isotopic labeling can be most effective for distinguishing the contributions of the limiting mechanisms.

## Introduction

Protein-protein association and dissociation are key to many cellular processes, ranging from transmembrane signaling^1–3^ to endocytosis. ^4^ While some protein complexes may involve little (re)structuring of the participating components and thus conform to a lock-and-key model of molecular recognition, it is now clear that often (un)folding and (un)binding are coupled.^5–7^ Coupled folding and binding can be described by two limiting mechanisms: induced fit, in which nonnative subunits form an initial encounter complex that then rearranges to a stable bound structure, and conformational selection, in which individual subunits first rearrange to conformations similar to those in the associated state and then bind. ^8^ Detailed characterizations of protein-protein association/dissociation simulations^9,10^ suggest that coupled folding and binding is often multipathway, combining elements of both of these limiting mechanisms. This makes both experimental and computational study of coupled folding and binding challenging.

The protein hormone insulin has come to serve as a model for studying coupled folding and binding owing to its small size and the therapeutic importance of its equilibrium between different oligomeric states.^11–14^ One such equilibrium is the one between dimer (Figure 1) and monomer. Each insulin monomer is 51 amino acids, organized into two polypeptide chains (A and B) joined by disulfide bonds (yellow in the left view). The 21-residue A chain forms two *α* helices (translucent), while the 30-residue B chain consists of an *α* helix (residues Ser^B9^-Cys^B19^, black) with a *β* turn (Gly^B20^-Gly^B23^, white) that leads to a C-terminal *β* sheet (Phe^B24^-Ala^B30^, red) in the dimer. Both experimental alanine scanning mutagenesis data^15^ and free energy simulations^16,17^ point to the importance of specific interfacial residues for stabilizing the dimer interface. These residues include the aromatic triplet of Phe^B24^-Phe^B25^-Tyr^B26^ on the interfacial *β* sheet,^13^ Tyr^B16^ on the interfacial *α* helix, and both Gly^B23^ and Pro^B28^ on the *β* turn and the C-terminal segment of the B chain, respectively.

**Figure 1:**
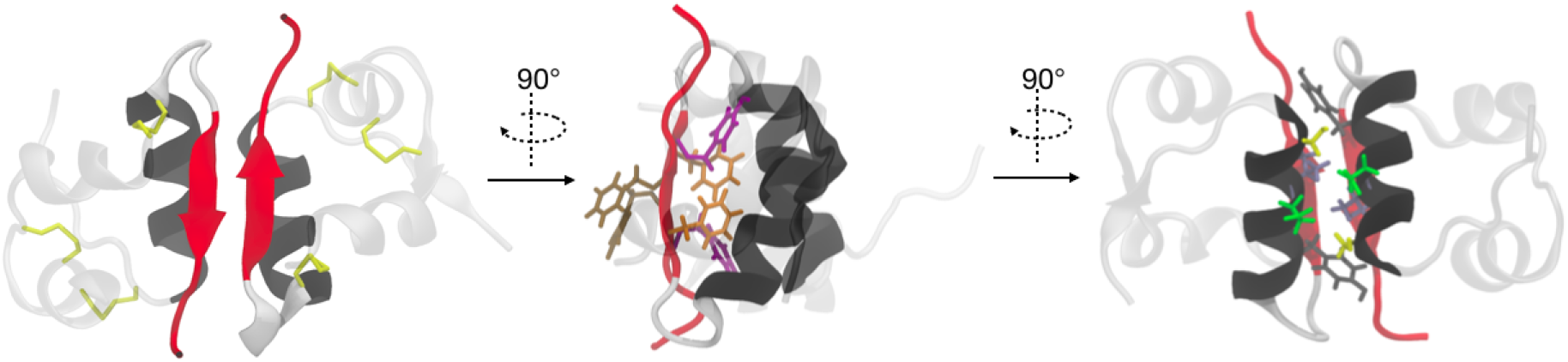
Three views of the insulin dimer. The A chain and residues Phe^B1^-Gly^B8^ of each monomer are shown in translucent silver, while interfacial residues are opaque. The interfacial *α* helices are shown in black, the *β* turn is shown in white, and the *β* sheet is shown in red. In the left panel, cysteine bonds are shown in yellow. In the middle panel, side chains for residues Phe^B24^ (orange), Phe^B25^ (brown), and Tyr^B26^ (purple) are shown. In the right panel, side chains for residues Ser^B9^ (yellow), Val^B12^ (blue), Glu^B13^ (green), and Tyr^B16^ (gray) are shown.

Dimer dissociation, which is a prerequisite for insulin to bind to its cellular receptor,^18^ is thought to be an example of coupled unfolding and unbinding. While the dimer is well structured,^19,20^ the monomeric state is thought to contain significant disorder. Specifically, experimental and computational studies indicate that Phe^B24^-Ala^B30^ can detach from the B-chain *α* helix and become at least partially disordered in the monomeric state.^19,21–26^ This detachment is thought to be important for insulin to bind its receptor^3,12,13^ based on structures of insulin in complex with fragments of the receptor.^27, 28^ An outstanding question is how Phe^B24^-Ala^B30^ detachment is coupled to dissociation of the dimer. More generally, the pathways of dimer dissociation remain poorly characterized. For example, it is unclear what role, if any, the interfacial *α* helices play in dissociation, and whether there are partially solvated or unfolded intermediates.

There is some experimental evidence suggesting the dimer dissociation could couple unfolding to unbinding. In particular, temperature jump two-dimensional (2D) amide-I infrared (IR) spectroscopy measurements suggest that, during dissociation, there is conformational rearrangement within the monomers on the timescale of 5 to 150 *µ*s, prior to loss of the *β* sheet at the dimer interface between 250 and 1000 *µ*s.^24,25^ Time-resolved X-ray scattering data also suggest an intermediate with conserved secondary structure on the timescale of 900 ns.^29^ These experiments, although mechanistically suggestive, provide limited structural information; complementary simulations are needed to microscopically interpret these data.

Recently, Bagchi and coworkers used metadynamics to compute the free energy as a function of the monomer-monomer center-of-mass distance and the number of intermolecular contacts, subject to restraints on the radii of gyration of the monomers.^30–32^ They identified a single major pathway of dissociation in which the number of intermolecular contacts was first observed to markedly decrease before the center-of mass distance increased. Through additional collective variables, they also characterized the protein-protein and protein-solvent interactions of Phe^B24^ and Tyr^B26^, indicating that conformational rearrangement and intra-monomeric unfolding are both coupled to the dissociation. Shaw and coworkers recently characterized the association of the insulin dimer through both unbiased simulation and tempered binding, an enhanced sampling technique that scales the protein interaction energies to encourage binding.^33^ In contrast to the simulations described immediately above, they found that successful association events consisted of insulin monomers adopting conformations similar to those found in the dimer before binding, and observed very little intramonomeric unfolding. The extent to which these two binding/unbinding pathways, one which involves monomeric unfolding and one which does not, can coexist is currently unknown.

In this work, we use a computational pipeline that combines multiple methods for enhanced sampling of rare events in molecular dynamics simulations to investigate coupled unfolding and unbinding during insulin dimer dissociation. In particular, we identify collective variables that fully resolve the possible pathways for the dissociation, and we show how an error estimator that we recently introduced^34,35^ can be used to quantitatively monitor convergence and allocate computational resources efficiently. The error estimator that we employ both provides quantitative evidence as to the convergence of our simulations, and allows us to meaningfully compare the free energy profiles of competing pathways by explicitly quantifying asymptotic errors. The computational pipeline enables us to show that there are multiple competing pathways for dimer dissociation, and we characterize these pathways in detail through additional collective variables that describe intra- and intermonomeric rearrangements.

The limiting behaviors observed correspond to induced fit and conformational selection mechanisms. Our simulations thus provide a unified perspective on the binding/unbinding paths observed in previous simulations. We go on to propose a set of experiments to investigate the relative contributions of our limiting paths. Specifically, we simulate IR spectroscopy experiments for a variety of isotope-labeled insulins and identify two labels which, when measured via T-jump IR spectroscopy, could experimentally distinguish the contributions from the limiting pathways to dimer dissociation. These simulated IR spectra also provide references to which future measurements can be compared, facilitating the interpretation of both equilibrium and T-jump IR spectra for the insulin dimer.

## Results and Discussion

Our goal was to investigate the molecular changes in intra- and intermolecular structure during insulin dimer dissociation. To this end, as detailed in Methods, we used steered molecular dynamics to generate multiple dissociation events and refined the resulting paths with the string method. Based on these simulations, we identified a small number of distances that provided good control over sampling (specifically, replica exchange umbrella sampling): 7 between C_*α*_ atoms in the interfacial *α* helices and 3 between C_*α*_ atoms in the interfacial *β* sheet (Figure 2). The *α* and *β* distances were separately averaged to define collective variables 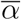 and 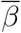, respectively. Below, we describe the potential of mean force (PMF) as a function of these coordinates, followed by additional statistical averages that provide further insights into specific intra- and intermolecular structural features. We conclude by showing how simulated vibrational spectra can serve as references for the design and interpretation of experiments.

**Figure 2:**
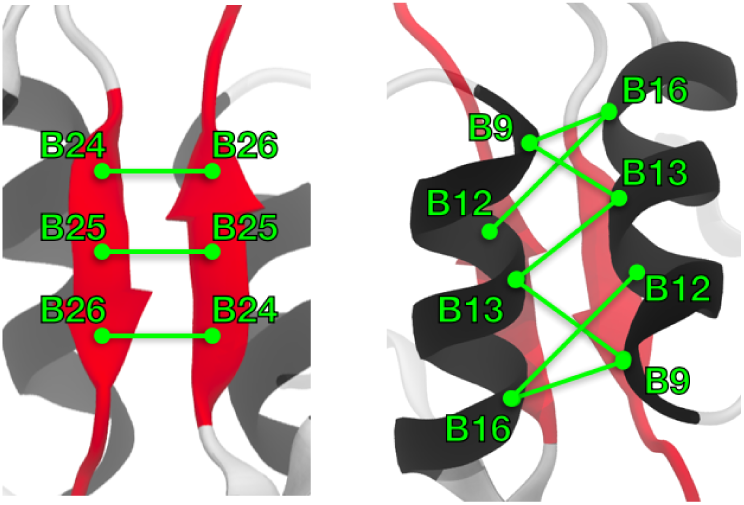
Schematic showing the *β* sheet contact pairs (left) and the *α* helix contact pairs (right). These correspond to the similarly labeled rows of Supplemental Table S1.

### Dissociation is multipathway, with two limiting cases

The PMF as a function of 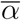 and 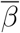 is shown in Figure 3, surrounded by representative structures. The minimum of the dimeric basin is marked by the circle at 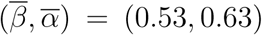 nm, and we set it to be the zero of free energy. The dimer is flanked by a trough along each axis, corresponding to breaking the *β* contacts while maintaining the *α* contacts and vice versa. There is a shoulder at 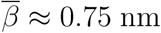; calculations described further below show that it coincides with solvent penetration of the *β* sheet. The remainder of the PMF is relatively flat. We take the monomeric state to be 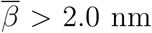 and 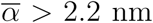 (marked by the dotted white box in Figure 3), which ensures that there are at least three layers of water between interfacial residues (see Methods). The free energy in this region ranges from 13 to 15.5 kcal/mol, within the range of previous estimates of the stability of the dimer.^16,17,30,36^ The plateau surrounding the monomeric state is between 1-3 *k*_*B*_*T* higher in free energy.

**Figure 3:**
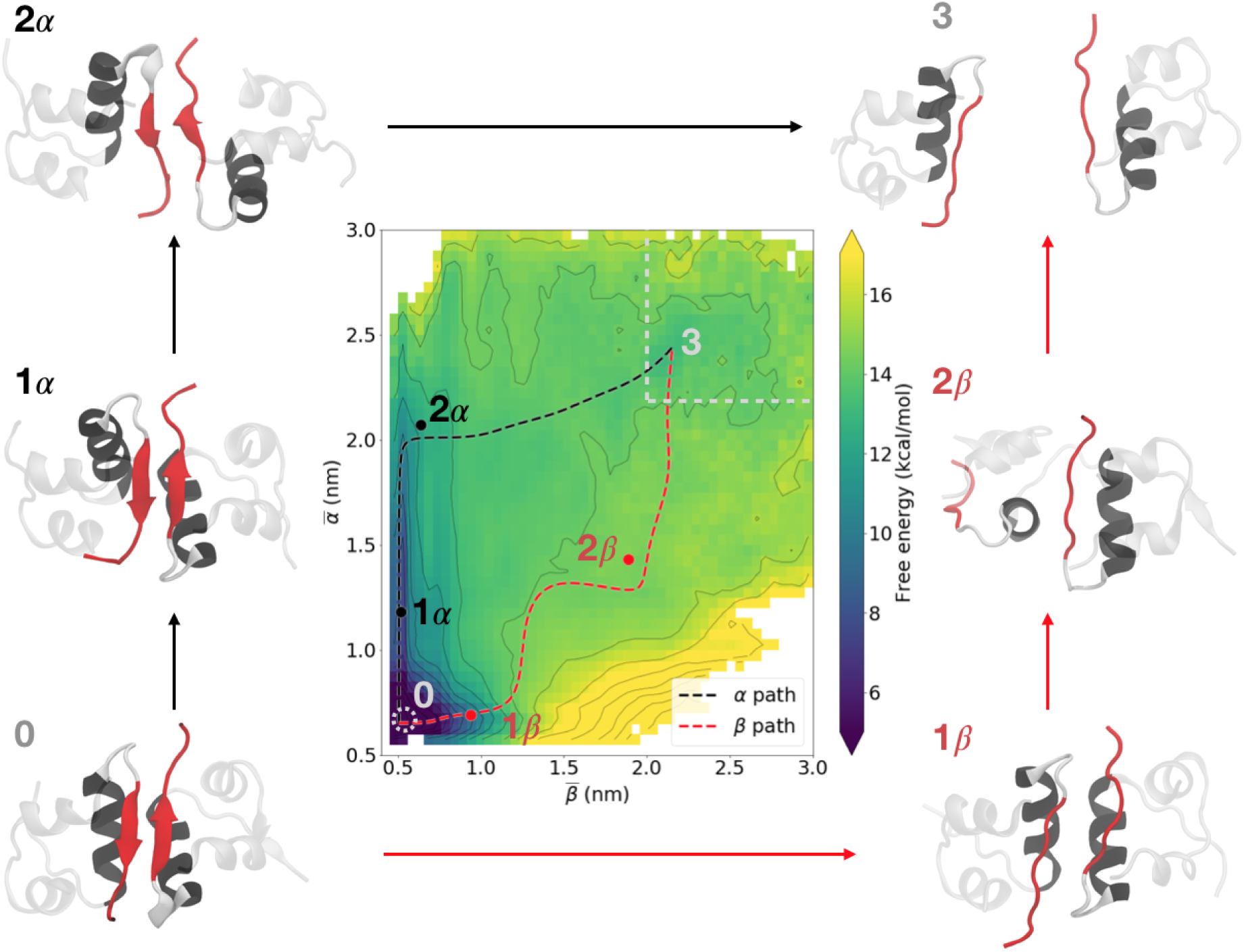
Potential of mean force (PMF) as a function of 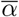 and 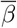. Limiting mean free energy paths in which the interfacial *α* or *β* contacts break first are indicated by black and red dashed lines, respectively. Representative structures corresponding to the marked points along the paths are labeled and shown adjacent to the PMF. These structures are referenced throughout the paper and are available in the supplemental material. The dimer is marked by a dotted white circle, and the monomeric state is marked by a dotted white box. Contour lines are every 2 *k*_*B*_*T*. The color scale is capped at both the upper and lower ends to more clearly show the variation in the partially-dissociated regime.

Consistent with the diversity of paths that we obtained in our steered molecular dynamics and string method simulations (see Methods), many minimum free energy paths can be drawn on the PMF (Supplemental Figure S1). These paths are stable not only in this 2D average distance space, but also the full 10-dimensional space of all individual distances, as indicated by the string method results in Supplemental Figure S1. Despite their similar maximum free energies, these paths imply dramatically different mechanisms of dissociation. For clarity, we focus on two limiting cases that initially follow the aforementioned troughs in free energy flanking the dimer. Along the *α* path (black in Figure 3), the interfacial *α* helices separate prior to the strands of the *β* sheet; along the *β* path (red in Figure 3), the order is reversed.

The free energy profile of the *α* path is consistent with the minimum free energy path obtained by Bagchi and coworkers (compare the black lines in Supplemental Figure S2 with Paths 1 and 4 in Figure 3 of ref. 30): there is an initial rise to an intermediate of 7.7 kcal/mol (5.3 kcal/mol in ref. 30), followed by a shoulder ∼2.4 kcal/mol higher in free energy and then a barrier of ∼4.0 kcal/mol. The free energy profile of the *β* path exhibits no comparable shoulders, but its maximum is comparable to that of the *α* path (14.7 kcal/mol and 14.1 kcal/mol, respectively). Considering the typical asymptotic variance of our PMF is on the order of 0.2 kcal^2^/mol^2^, we thus expect both of these limiting paths, as well as the many paths that fall between them (Supplemental Figure S1), to contribute to dissociation.

### The monomers rotate relative to each other in opposite ways along the two limiting paths

Previous studies ^30,31^ reported rotation of the interfacial *β* strands relative to each other, as characterized by a pseudodihedral angle (here denoted Φ_*β*_) defined by the C_*α*_ atoms of Tyr^B26^, Phe^B24^, Tyr^B*′*26^, and Phe^B*′*24^, where the primes distinguish one monomer from the other. In addition to Φ_*β*_, we calculate the pseudodihedral angle (Φ_*α*_) between the geometric centers of the backbone atoms of the residues that define the dimeric interfacial alpha helices: Ser^B9^-Leu^B11^, Leu^B17^-Cys^B19^, Leu^B*′*17^-Cys^B*′*19^, and Ser^B*′*9^-Leu^B*′*11^. Explicit illustrations of these angles are shown in Supplemental Figure S3. By examining both Φ_*β*_ and Φ_*α*_, we can better understand whether the rotation is restricted to the *β* strands, or if the entire dimer interface moves together. Since molecular dynamics simulations allow any collective variable to be calculated within machine precision, we create PMFs in four 2D spaces that measure interfacial rotations as both 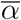 and 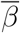 change (Figure 4A); the plots are restricted to ranges of 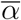 and 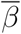 that correspond to the initial steps of dissociation because the interfacial pseudodihedrals become poorly defined when the average distances are large. All of the PMFs, both in Figure 3 and Figure 4, are generated from the same dataset, as EMUS allows for the calculation of PMFs in arbitrary collective variable spaces without the need for additional sampling.

**Figure 4:**
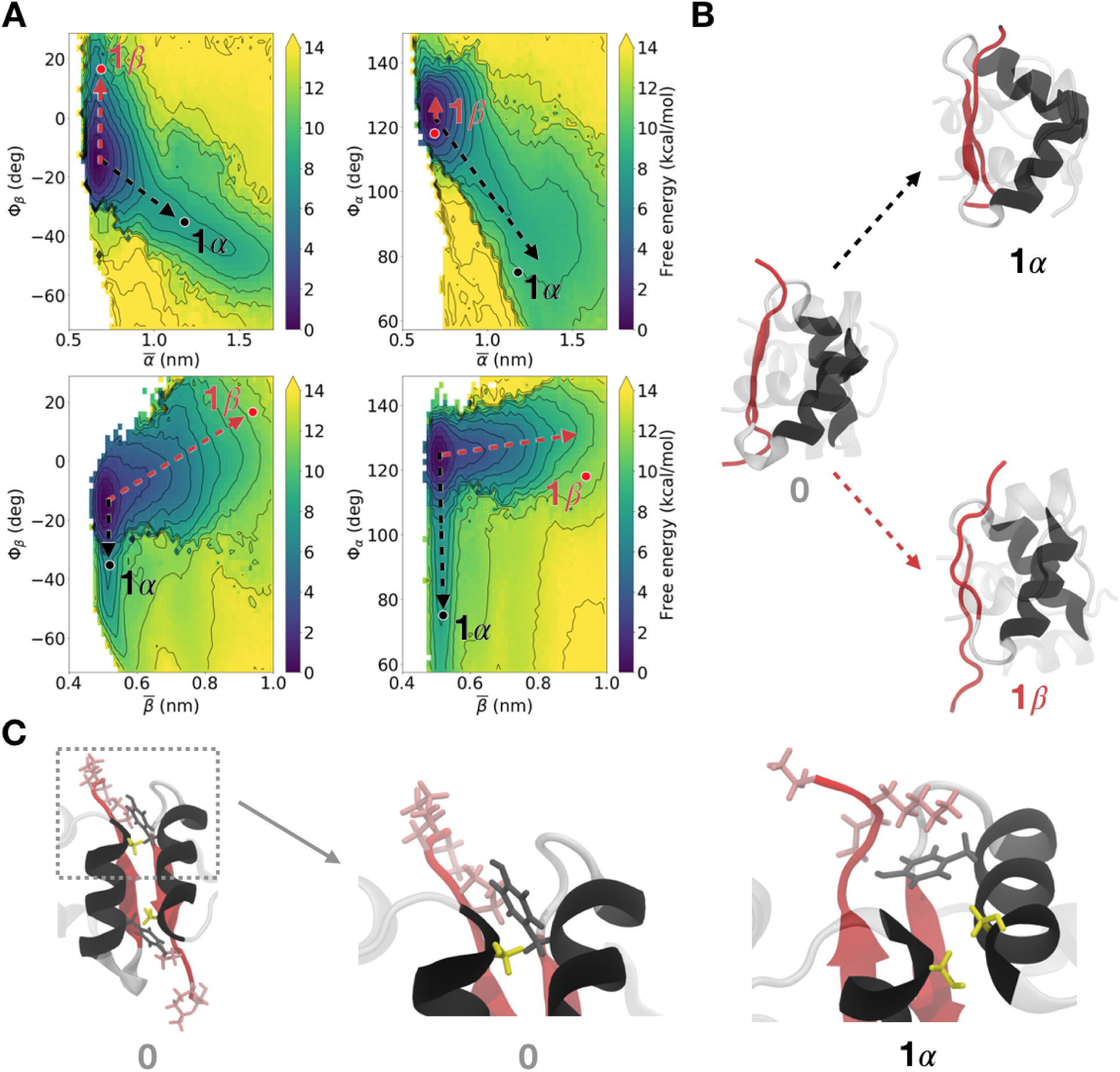
The monomers rotate relative to each other during dissociation. (A) PMFs characterizing the rotations as pairwise functions of 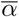 (top) or 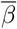 (bottom), and Φ_*β*_ (left) or Φ_*α*_ (right). Superimposed arrows show the negative rotations associated with the *α* path (black) and the positive rotations associated with the *β* path (red). Intermediates are marked on the PMF, and are as labeled in Figure 3. Structures were chosen to show the rotation of Φ_*β*_; for this reason, the arrows in the left plots terminate at the dots but those on the right plots do not. Contour lines are every 2 *k*_*B*_*T*. The color scale was capped at 14 kcal/mol. (B) Representative structures for the rotations along the *α* and *β* paths, represented by the black and red arrows, respectively. These structures, labeled in (A), are the same as those labeled in Figure 3. (C) The dimer with the interfacial *α* helices in front, showing the side chains for Ser^B9^, Tyr^B16^ (gray) and Pro^B28^-Ala^B30^ (pink). Zooming in (middle), one can see the native contact of Ser^B9^-Tyr^B*′* 16^, with Pro^B28^-Ala^B30^ behind. Along the *α* path (right), Tyr^B*′* 16^ has rotated away from Ser^B9^, and is instead in contact with Pro^B28^-Ala^B30^. Furthermore, this rotation brings Ser^B9^ and Ser^B^*′*^9^ together.

There is a deep free energy basin corresponding to the dimer at 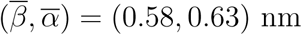 and (Φ_*β*_, Φ_*α*_) = (−15*°*, 125*°*). This reflects the fact that in the dimer state, in agreement with available crystal structures, there is a slight rotation from parallel between the *β* sheet residues, and a more pronounced rotation between *α* helices, consistent with well-known characterizations of *α* helix packing.^37–39^ The PMFs are dominated by the troughs flanking the dimer in Figure 3. The trough along the *α* path (black dotted arrows) is readily visible in both sets of plots and shows that increases in 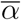 are coupled to negative rotations of both Φ_*α*_ and Φ_*β*_ (by −35*°* and −45^*°*^, respectively). The trough along the *β* path (red dotted arrows) is more readily visible in the bottom plots and shows that increases in 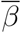 are coupled to positive rotations of Φ_*β*_; there is little change in Φ_*α*_, suggesting the interfacial *α* helical contacts are maintained. Side views (similar to the middle panel of Figure 1) of the dimer and rotated species show the structural consequences of these interfacial rotations (Figure 4B). Namely, comparing these side views with the corresponding front views (Figure 3) suggests that the initial dissociation along the limiting *β* path involves the breaking of the interfacial *β* sheet and the positive rotation of Φ_*β*_. In contrast, the initial dissociation along the *α* path comes as the *α* helices twist away from each other, coupled with negative rotations of both Φ_*α*_ and Φ_*β*_.

The negative rotations along the *α* path enable formation of nonnative interactions (Figure 4C). The serine side chains of Ser^B9^ and Ser^B*′*9^ (yellow in Figure 4C) form a hydrogen bond at 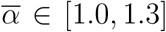 nm. Then, as the *β* strands rotate relative to each other along the *α* path, Pro^B28^-Ala^B30^ (pink) breaks its native contacts and instead forms a contact with tyrosine Tyr^B*′*16^ (gray), in the *α* helix of the opposite monomer. This tyrosine at Tyr^B16^ contacts Ser^B9^, Val^B12^, and Tyr^B*′*26^ in the dimeric state (see Supplemental Table S1), but these interactions are broken as the *α* helices separate. Averages of side chain contacts that quantitatively show these trends are seen in Supplemental Figure S4. Furthermore, the necessity of breaking the native contacts between Pro^B28^-Ala^B30^ and Gly^B*′*20^-Gly^B*′*23^ as the dissociation progresses is consistent with the mutations that yield fast-acting insulin analogs, discussed further in the Supplemental Information (Supplemental Figure S5). No comparable nonnative interactions are observed along the *β* path.

These projections further allow us to compare with Bagchi and coworkers’ results. In particular, Figure 8 of ref. 31 indicates a path which begins with a slight increase in Φ_*β*_ coupled to an increase in 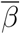 from approximately 0.6 to 1.1 nm. This initial step is followed by a 30*°* decrease in Φ_*β*_ coupled to a return to a dimer-like 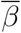 (near 0.6 nm) as the dissociation progresses. The *β* strands only separate at the final step of their described mechanism. We interpret this to correspond to taking an initial step along the *β* path, then collapsing back to a near-dimer like *β* interface before proceeding along the *α* path. That said, the projections of the *α* and *β* paths onto Bagchi and coworkers’ coordinates do not fall precisely on top of their minimum free energy path (Supplemental Figure S6). By explicitly probing the multipathway nature of the dissociation, our results reveal the homogeneity of rotation profiles depending on dissociation path.

### Water solvates key interfacial residues as dissociation progresses

Solvent plays a key role in protein association/dissociation processes. Moreover, time resolved X-ray scattering data suggest at least one intermediate in the insulin dimer dissociation that involves quick solvent uptake by a species with dimer-like secondary structure, coincident with a slight increase in molecular volume.^29^ To investigate the possible presence of a similar feature in our simulations, we defined three collective variables; in order of increasing specificity, they are (i) the total molecular volume, (ii) the solvent accessible surface area (SASA) of eight residues that make up the hydrophobic core of the interface (Val^B12^, Tyr^B16^, Phe^B24^, Tyr^B26^ on each monomer), and (iii) the number of native interfacial hydrogen bonds, which only form between Phe^B24^ and Tyr^B26^. The total molecular volume, probed by the X-ray scattering, can reflect solvent uptake, but it can also correspond to large-scale conformational change. The core SASA reflects the solvation of the interface in general, while the hydrogen bonding between interfacial residues probes the loss of dimer-like secondary structure. Averages of these variables are seen in Figure 5A.

**Figure 5:**
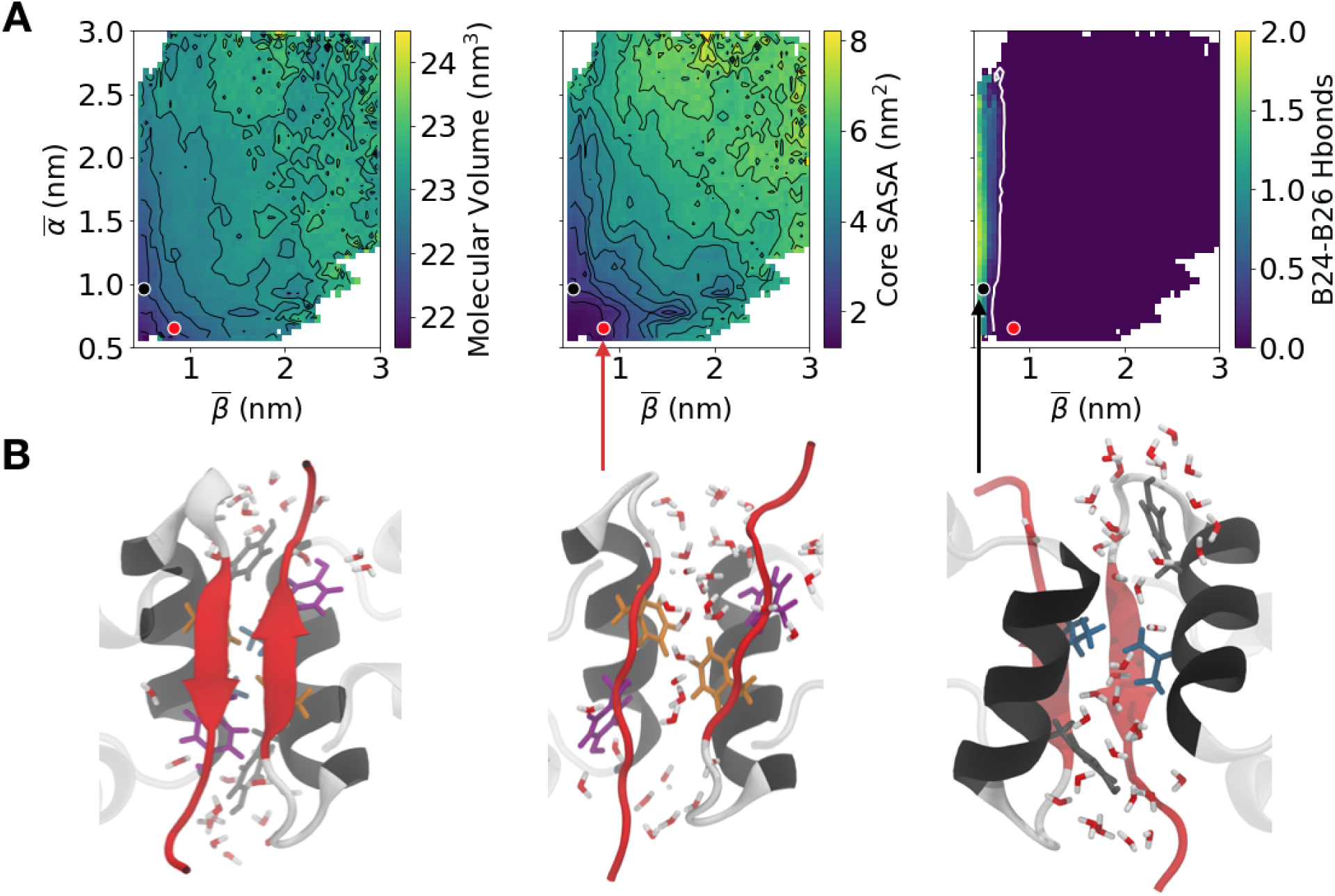
Characterizing solvation. (A) Averages of total molecular volume (left), core SASA (middle), and number of interfacial Phe^B24^-Tyr^B26^ hydrogen bonds (right) as a function of 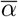 and 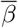. Contours are every 0.2 nm^3^ and 0.5 nm^2^ for the molecular volume and SASA plots, respectively. The white contour on the right plot indicates where the number of hydrogen bonds drops to 2% of the average in the dimer. (B) Insulin structures showing the unsolvated dimer interface (left), the solvation of the *β* interface (middle), and the solvation of the *α* interface (right). The locations of these structures are marked in (A).

The red and red black dots in Figure 5A mark the positions of the structures in Figure 5B, which show characteristic solvations of the *β* and *α* interfaces, respectively. These structures represent low free energy states relative to the barriers along the *β* or *α* paths. Moving from the dimeric state to either of these positions, the total molecular volume increases by 0.3-0.4 nm^3^. This small increase in molecular volume allows some solvation of the interface, increasing core SASA by between 0.8-1.2 nm^2^. Along the *α* path, this solvation is at the *α* interface, while, along the *β* path, this solvation is at the *β* interface (Figure 5B). The former does not result in a loss of secondary structure as measured by STRIDE.^40^ By contrast, we do see a distinct loss of *β* sheet content along the *β* path. Specifically, as 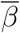 increases, the Phe^B24^ and Tyr^B26^ hydrogen bonds across the interface are replaced with ones to solvent (white contour in the right panel in Figure 5A and Supplemental Figure S7), signaling the loss of the interfacial *β* sheet.

One would expect that the loss of the interfacial *β* sheet and the solvation of the hydrophobic core are highly correlated because the separation of the *β* strands allows water to penetrate between the monomeric units. We observe this to be the case when the *α* helices are already partly separated 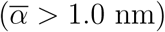. The breaking of the interfacial hydrogen bonds, represented by the white contour in the right panel of Figure 5A, occurs in the same area of the collective variable space as the rapid solvation of the hydrophobic core, represented by the tight black contours in the middle panel of Figure 5A. This area, where 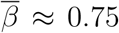 nm, is also co-located with the shoulder in the PMF mentioned earlier (Figure 3), suggesting that the loss of *β* sheet content and concomitant solvation gives rise to a slight decrease in free energy. The stabilization that we observe is consistent with previous simulations of mutants of the insulin dimer which indicate that water can mediate *β* sheet interactions by forming hydrogen bonds that bridge between residues.^36^ The existence of this shoulder is also consistent with states involved in the dewetting transition seen in ref. 32, in which the center-of-mass separation of the monomers is 2 nm and there are a few water molecules at the interface.

However, when the *α* helices are in a near-native distance 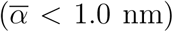, the solvation of the hydrophobic core occurs after the loss of the interfacial *β* sheet. In this case, the protein-protein hydrogen bonds are broken when 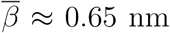 (white contour in the right panel of Figure 5A) but the hydrophobic core residues do not become significantly solvated until the *β* strands are even further separated, at approximately 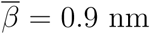 (middle panel of Figure 5A). This occurs within a low-free-energy trough of the PMF; in other words, the PMF only rises sharply as the core is solvated, not with the loss of hydrogen bonds. Evidence of these dynamics are further seen in the simulated IR results presented later, in which the peak absorption/emission doublet is red shifted with only a minimal loss in intensity.

As mentioned in the introduction, Chen and coworkers found evidence for a partially-solvated intermediate with dimer-like secondary structure, implying conserved interfacial *β* sheet character.^29^ Although we find no distinct free energy basin that obviously corresponds to this intermediate, the initial partially solvated structures we observe along the *α* path are consistent with these data, as the interfacial *β* sheet is conserved. However, even with the loss of the *β* sheet along the *β* path, our simulations do not rule out the possibility of structures along the *β* path also being consistent with the X-ray scattering data. Specifically, the experimental data were collected at 316 K and 0.27 M DCl (pH ≈ 0) in a solution of ethanol and water, while our simulations are for 303.15 K and pH 7 with only water as the solvent. Previous computational work suggests that ethanol appears to facilitate partial solvation of the *β* interface,^31^ so further simulations are needed to definitively interpret these T-jump X-ray scattering experiments.

### The B-chain C-terminal segment detaches along the *α* path but not the *β* path

As noted above, there is extensive evidence that, for insulin to bind its receptor, the B-chain C-terminal segment must detach from the B-chain *α* helix.^13,27,28,41,42^ Detachment and partial unfolding have also been invoked to explain the diagonal elongation of features in equilibrium 2D infrared spectra of insulin at elevated temperatures. ^24^ Whether detachment and partial unfolding occurs during dissociation remains an open question.

These considerations, combined with the known partial disorder of the *β* turn and the B-chain C-terminal segment in the insulin monomer,^22–24,26^ motivated our definition of two average angles to study intramonomeric unfolding during dimer dissociation: 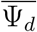, which measures detachment of the B-chain C-terminal segment from the B-chain *α* helix, and 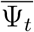, which characterizes the disorder of the *β* turn. Results for 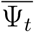 are discussed in the Supplemental Information (Supplemental Figure S8); here we focus on 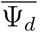. 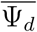 is the angle between the C_*α*_ atoms of Arg^B22^, Phe^B24^, and Tyr^B26^ (Supplemental Figure S9), measured for each monomeric unit and then averaged. 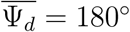 indicates an attached structure, because a flat *β* strand tucks against the B-chain *α* helix. In contrast, 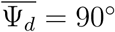 indicates a structure that is almost completely detached, with the B-chain C-terminal segment bent away from the *α* helix. This detachment is also coupled to the solvation of Gly^A1^-Val^A3^, which are involved in binding to the insulin receptor^20^ (Supplemental Figure S10). Examples of monomeric structures with attached and detached B-chain C-terminal segments are shown in Figure 6A. Figure 6B shows the full dimer view of the same detached intermediate with Ψ_*d*_ = 140*°* to illustrate how the detachment of the C-terminal segment allows for the nonnative interaction of Pro^B28^-Ala^B30^ and Tyr^B*′*16^, the same nonnative interaction shown in Figure 4C.

**Figure 6:**
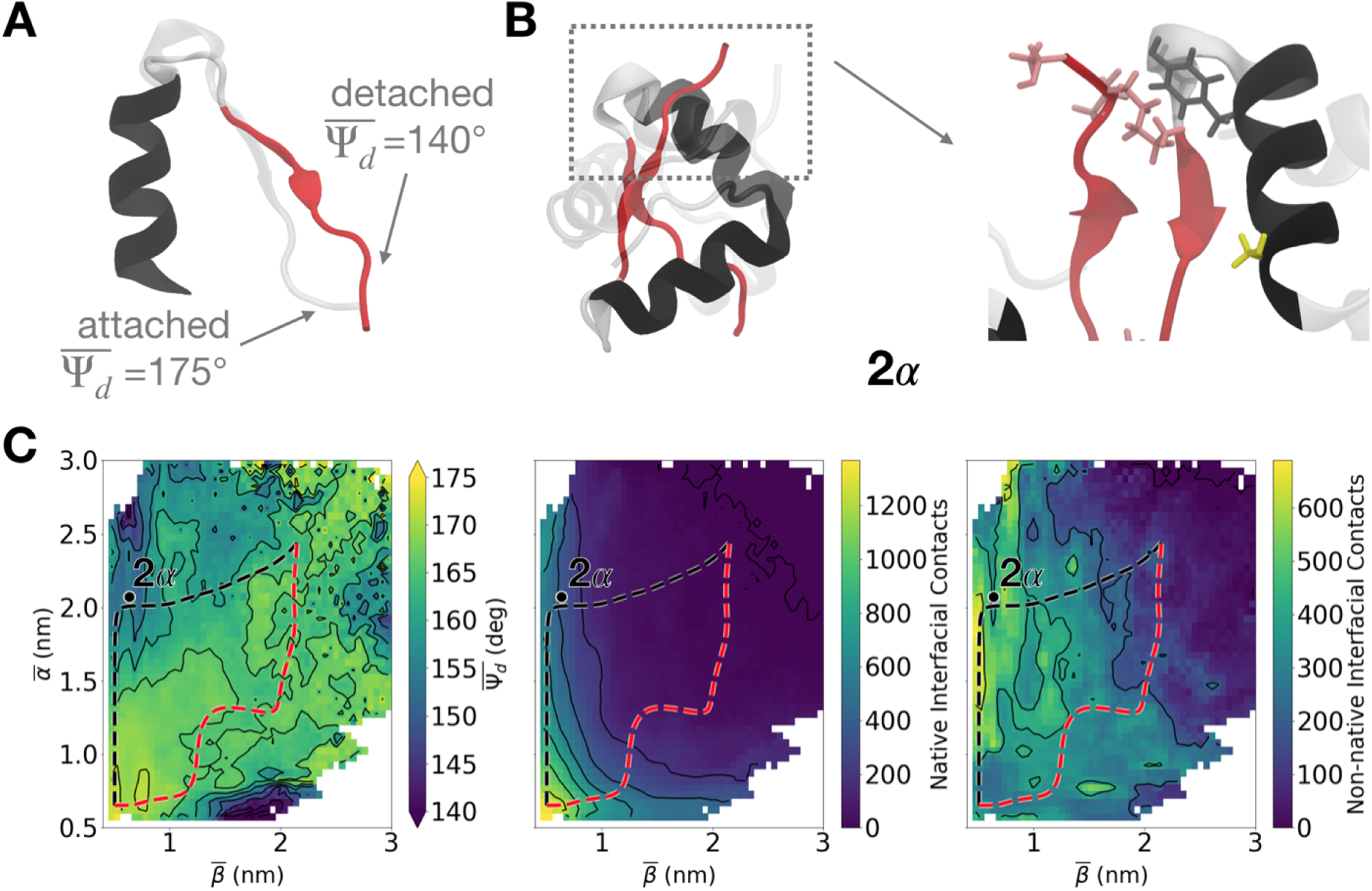
Characterizing detachment. (A) A representative monomeric structure contrasting attached and detached B-chain C-terminal segments. (B) Structural depiction of how the detachment of the B-chain C-terminal segment allows for continued nonnative interactions between Pro^B28^-Ala^B30^ and Tyr^B*′* 16^. (C) (Left) Average of 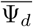 as a function of 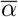 and 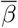 with black contour lines shown every 5*°*. (Middle) Number of native non-hydrogen atom native interfacial contacts and (right) non-hydrogen atom nonnative interfacial contacts (cutoff 7 Å), with contour lines shown every 200 contacts. On all graphs, the *α* (black) and *β* (red) paths are shown, as is the location of structure 2*α* shown in (B).

The average value of Ψ_*d*_ as a function of 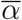 and 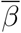is plotted in the left panel of Figure 6C. The dimer state corresponds to 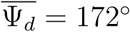, consistent with the *β* strand being attached. 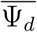 decreases significantly along the *α* path; the latter half of the path 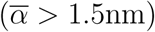 consists mainly of structures in which the B-chain C-terminal segment is detached 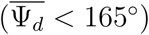. This behavior contrasts with the *β* path, in which the minimum value of 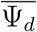 is 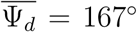 (we neglect the low values of 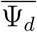 in the lower right corner of Figure 6C because that region is very high in free energy; see Figure 3). Overall, while we observe some detachment along the *β* path, it is much more pronounced along the *α* path. It is also worth noting that in the monomeric region identified earlier (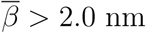 and 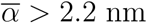) the detachment is less pronounced than at intermediate stages of the *α* path. However, the monomeric region has more variability in detachment angle than in the dimeric region, consistent with previous results showing a limited amount of disorder in the C-terminal segment of the B chain.^22–24,26^

Along the *β* path, instead of unfolding, the *β* sheets separate and the monomers drift away from one another, forming a diverse set of non-specific, nonnative interfacial contacts, as in structure 2*β* in Figure 3. The numbers of native and nonnative interfacial contacts are plotted in the center and right panels of Figure 6C, respectively. Along the *β* path (red), the native contacts are almost completely broken as 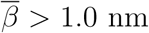, although a limited number of nonnative contacts persist as the dissociation proceeds further. Similarly, along the *α* path (black), we also see the formation of nonnative contacts coupled to the breaking of native contacts, consistent with the side-chain interactions discussed previously (Figure 6B).

Finally, we note that T-jump 2D amide-I IR spectroscopy experiments in 20% ethanol indicated two contributions to the dimer dissociation process: melting of the dimer *β* sheet observed between 250-1000 *µ*s, and a 5-150 *µ*s process that was assigned to *α* helix disordering.^25^ These timescales thus suggest that unfolding occurs before the loss of the interfacial *β* sheet. In our (aqueous) simulations, we do not observe any loss of *α* helix content, although it may be possible that helix rotation could give rise to such a signal. Instead, the unfolding that we observe is restricted to detachment of the B-chain C-terminal *β* strand and disorder of the *β* turn (see Supplemental Figure S8), which primarily occurs along the *α* path. Furthermore, the detachment along the *α* path (following the red path in the left panel of Figure 6C) starts to occur before the loss of the interfacial *β* sheet (right panel of Figure 5A). The *α* path, which exhibits monomeric unfolding in the form of C-terminal detachment while maintaining dimer-like secondary structure, is thus consistent with the T-jump data gathered by Tokmakoff and coworkers.^25^ Moreover, this detached state also corresponds to an increase in molecular volume of between 0.6 and 0.8 nm^3^, which is consistent with the evidence from Chen and coworkers for a second intermediate that corresponds to a large increase in molecular volume while maintaining secondary structure.^29^

### The *α* and *β* path correspond to induced fit and conformational selection

In this section, we connect the intramonomeric detachment and core solvation with the inter-monomeric rotations discussed earlier to characterize the various mechanisms of the insulin dimer dissociation, making explicit comparisons to the induced fit and conformational selection models of coupled folding and binding.

The *α* path, as mentioned before, initially consists of rotations at both the *α* and *β* interfaces (Figure 4A); the alpha helices twist away from each other, forming nonnative contacts between Ser^B9^:Ser^B*′*9^ and Pro^B28^-Ala^B30^:Tyr^B*′*16^ (Figure 4A). This initial *α* helix separation and rotation increases the SASA of the hydrophobic core by ∼1.5 nm^2^, while further *α* helix separation (between 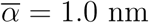 and 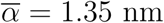) leads to very little additional core solvation (Figure 5A). This solvation correlates with the free energy along the *α* path, which also sharply increases until 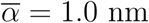, where it levels off around 8 kcal/mol until 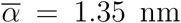 (Figure 3). There is then another free energy barrier of 2 kcal/mol when 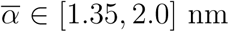, which correlates well with the *β* sheet starting to detach from the core (Figure 6C). This detachment, while partially maintaining the nonnative interactions between Pro^B28^-Ala^B30^:Tyr^B*′*16^, sacrifices native contacts and exposes Gly^A1^-Val^A3^ to the solvent (Supplemental Figure S10). The last barrier of 4 kcal/mol correlates to the breaking of the interfacial *β* sheet hydrogen bonds and the subsequent separation of the *β* strands. Structurally this final step is heterogeneous, involving varying amounts of detachment, rotation, and sliding of the *β* strands as 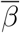 increases.

When thinking of the association process, we posit that the *α* path corresponds to the induced fit model of coupled folding and binding. Following the black trace in Figure 6C from monomer to dimer (top right to bottom left), association is initiated by the B-chain C-terminal strands detaching and encountering one another. The *β* interface becomes structured; then, the *α* helices are recruited into the interface, leading to a dimer-like structure. The monomers only adopt their structures in the dimer upon association.

The *β* path first couples the initial separation of the *β* strands to their rotation (Figure 4A). This correlates to the sudden increase in free energy from 0 to 14 kcal/mol as 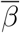 increases from 0.5 to 1.2 nm. The subsequent steps of the dissociation correspond to the two monomers, with conformations similar to the ones found in the dimer, drifting away from one another. Specifically, as the dissociation proceeds, a diverse set of nonnative interfaces are formed that are structurally close to the native interface, but involve minimal detachment of the B-chain C-terminal segment (Figure 6C). This is consistent with the relatively flat free energy trace when 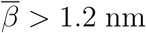 (Supplemental Figure S2), as all of these near-native, folded structures are similar in terms of solvation and protein-protein interactions.

Thinking of the association process for this limiting path, we posit that the *β* path corresponds to the conformational selection model of coupled folding and binding. Following the red trace in Figure 6C from monomer to dimer, the monomers first adopt structures like those in the dimer, with attached B-chain C-terminal segments; they then approach one another, forming a variety of nonnative interfacial contacts that eventually collapse to the dimer-like set of native contacts. We note that the *β* path is consistent with the unbiased association trajectories simulated by Shaw and coworkers, who found that insulin dimer association is characterized by monomers, largely folded into their conformations in the dimer, forming near-native interfaces that eventually collapse to the native dimer interface.^33^ In contrast, the biased association trajectories from ref. 33 primarily exhibit interfacial rotations and nonnative interactions (namely, Ser^B9^:Ser^B*′*9^) consistent with the *α* path, albeit without substantial B-chain C-terminal detachment. To remind the reader, the free energy profile, interfacial rotations, and monomeric unfolding observed along the *α* path are all consistent with the free energy minimum path discussed by Bagchi and coworkers. ^30^ The collective variables we identified enabled us to obtain an ensemble of paths that encompassed the diversity seen in previous studies.^30,33^

### Simulations enable determination of optimal isotopic labeling sites for infrared spectroscopy

Even though the *α* and *β* paths are only limiting cases, most of the paths in Supplemental Figure S1 initially follow one of these limiting paths. As discussed previously, the initial steps along the *α* and *β* paths correspond to solvation of the *α* and *β* interfaces, respectively. In particular, the *α* path consists first of the solvation of the *α* interface, followed by the solvation of the *β* interface; the *β* path reverses this ordering. Thus, these simulation results can be further investigated by experimental techniques that can resolve residue-level solvation.

Fourier-transform (FT) and two-dimensional (2D) amide-I infrared (IR) spectroscopies are useful for studying protein secondary structure and solvation because they are sensitive probes of the hydrogen bonding of the carbonyl groups in protein backbones. In particular, one typical hydrogen bond to an amide carbonyl causes a redshift of about 16 cm^*−*1^.^43^ This means the location of the carbonyl stretch is sensitive to the number and strength of hydrogen bonds made by the backbone carbonyls, which includes hydrogen bonds associated with both secondary structure and protein-solvent interactions. Moreover, one can isotopically label specific amide carbonyl groups with ^13^C^18^O, redshifting their vibrations by 65 cm^*−*1^ to isolate them from the other amide vibrations. Both 2D and 1D IR spectra can be generated through molecular modeling, effectively “mapping” the classical variables from molecular trajectories into a quantum-mechanical Hamiltonian (see ref. 43 for further review of 2DIR methods and their simulation).

These simulated spectra have been used to interpret both equilibrium and T-jump measurements of protein folding. ^44^ Furthermore, recent spectral simulation work from Meuwly and coworkers has shown that isotope labeled spectra for both the insulin dimer and monomer are qualitatively sensitive to the number of waters hydrating the labeled backbone carbonyl group.^45^ They note, however, that no one structural feature is particularly strongly correlated to spectroscopic behavior, which is instead sensitive to the rapidly fluctuating environment around each backbone oscillator. When considering how best to experimentally probe the dynamics of the dissociation, it is thus difficult to *a priori* suggest the best sites to label. In previous sections, we discussed our results in the context of previous equilibrium^24^ and T-jump^25^ experiments of unlabeled insulin. Below, we combine our umbrella sampling results with additional simulations of IR spectra to propose new sites for isotopic labeling, with a view toward achieving residue-specific characterization of the dimer dissociation mechanism.

We expected the most promising sites for isotopic labeling to be at the dimer interface, as the participating residues exhibit large changes in SASA upon dissociation (Figure 5). Owing to the computational cost of 2DIR simulations, we first simulated FTIR spectra along both limiting paths for all possible constructs with a single interfacial residue isotopically labeled (between Ser^B9^ and Ala^B30^). This process, summarized in the Supplemental Information (see Supplemental Figure S11), revealed that the isotopic labels that produce simulated spectra most sensitive to solvation are those on the residues in the previously identified hydrophobic core. For clarity, we focus on two specific residues: Phe^B24^, at the *β* interface, and Glu^B13^, at the *α* interface. A construct with the former label was studied in refs. 45 and 46, while, to the best of our knowledge, a construct with the latter label was not previously synthesized. For each label, we identify and simulate 2DIR for three states: the dimeric state, the monomeric state, and the solvated state (Figure 7). The solvated states for the Phe^B24^ and Glu^B13^ labels represent partially dissociated species with solvated *β* and *α* interfaces, respectively. The process of defining these solvated states is described in the Supplemental Information (Supplemental Figure S11).

**Figure 7:**
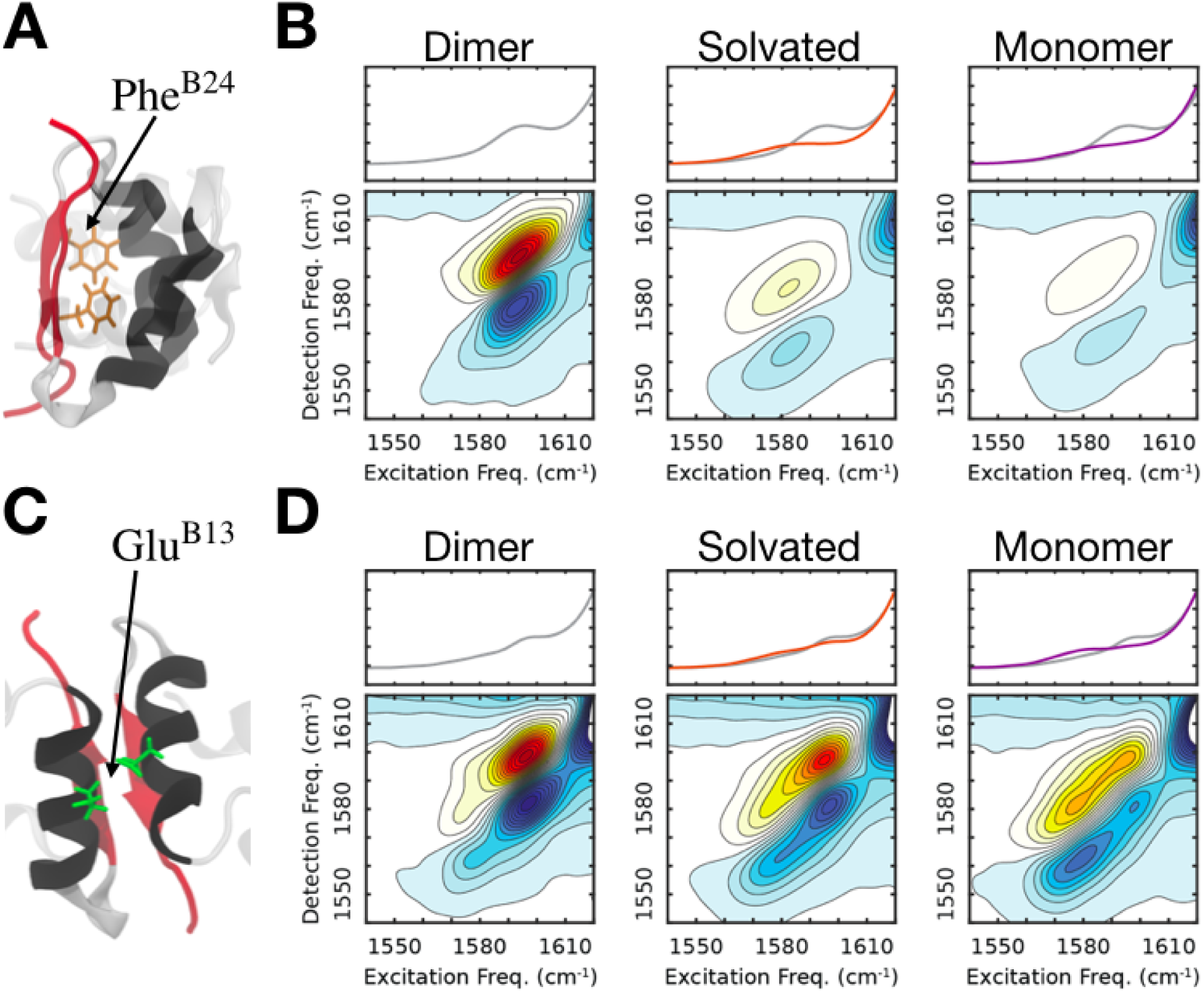
Simulated IR spectra for selected isotopically labeled constructs. (A) Dimeric structure showing the Phe^B24^ side chain, which was isotopically labeled on its backbone carbonyl. (B) Simulated 2DIR spectra of the Phe^B24^-labeled dimer (left), solvated species (middle), and monomer (right). Intensities are normalized using the peak intensity of the dimer spectrum, with the contours spaced by 7.5%. (C & D) Similar structures/spectra, but for the Glu^B13^ -labeled insulin. In both cases, the spectra for the solvated species were generated from structures along both the *α* and *β* paths.

**Figure 8:**
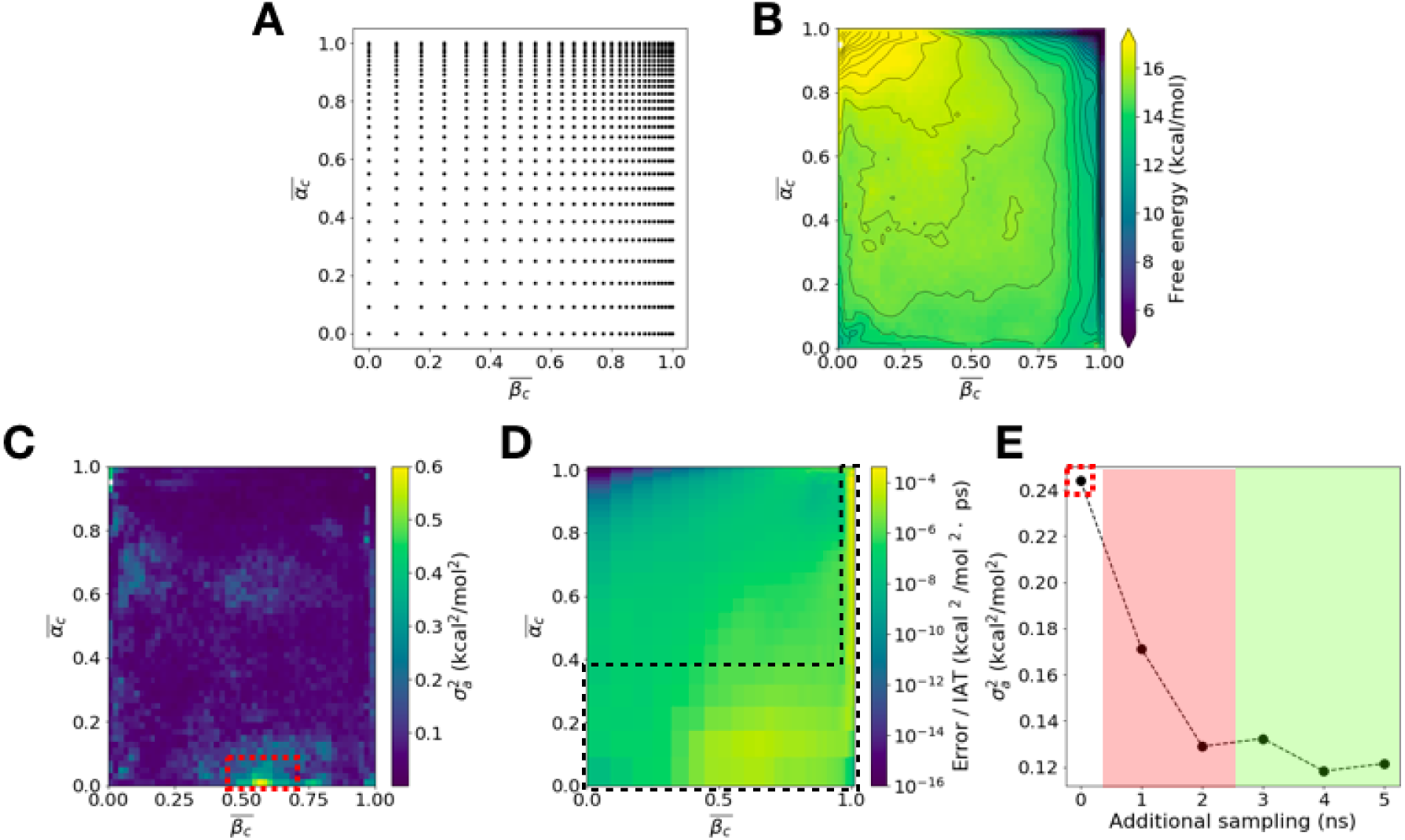
Umbrella sampling. (A) The location of the window centers used for the REUS procedure, shown in the space of 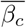 and 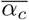. These are logarithmically spaced to place more density near the dimer (upper right corner). (B) Free energy as a function of the average numbers of *β* and *α* contacts at the insulin dimer interface (contour spacing 0.5 kcal/mol). 5 ns of sampling was gathered per window (784 windows). (C) The asymptotic variance associated with the free energy in (B). The region of highest variance, with average and maximum 0.59 kcal^2^/mol^2^, is marked by the red box. (D) The per-window error contributions to the marked variance in (C), assuming that the matrix Σ is diagonal. 5 ns of additional sampling was added to only the boxed black area of large error contributions. (E) How the average asymptotic variance of the marked region in (C) decreased as 5 more ns of sampling was added per selected window. The red shaded region represents the area where the additional sampling is shorter than 10 times the autocorrelation time for EMUS quantities. The asymptotic variance data in this region is thus unreliable. Reliable asymptotic variances are obtained in the green shaded region.

For insulin labeled at Phe^B24^ (at the *β* interface), there is a strong peak at 1595 cm^*−*1^ in the dimer spectrum. This feature decreases in intensity, becomes redshifted to 1582 cm^*−*1^, and broadens significantly along the diagonal as the carbonyl group of Phe^B24^ becomes solvated, consistent with previous measurements on the monomer.^46^ Microscopically, we interpret the changes to reflect the hydrogen bond between Phe^B24^ and Tyr^B*′*26^ breaking and the backbone becoming solvated. This behavior was observed once 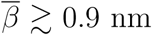 along both the *α* and *β* paths (Supplemental Figure S11), suggesting that 2DIR experiments using this label should serve as a probe for the solvation of the *β* interface regardless of the dissociation mechanism. For most regions of the collective variable space, the redshifting and broadening of the peak occur together, but along the *β* path at 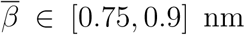, the redshifting precedes the broadening. This corresponds to the interfacial hydrogen bonds breaking prior to solvation of the Phe^B24^ backbone carbonyl (Figure 5).

Similarly, for insulin labeled at Glu^B13^ (at the *α* interface), we see a peak at 1595 cm^*−*1^, and it becomes redshifted to 1579 cm^*−*1^ and broadens along the diagonal as the site containing the label becomes solvated. Again, this behavior was observed along both the *α* and the *β* paths (Supplemental Figure S11), so 2DIR experiments of this construct should serve as a probe for the solvation of the *α* interface regardless of the dissociation mechanism. By using the Phe^B24^ and Glu^B13^ isotopic labels in T-jump experiments, one should be able to determine the times that the *α* and *β* interfaces become solvated and in turn the order of events during dissociation. In this way, one might tell if one limiting mechanism or the other predominates.

## Conclusions

A key step in insulin function is its dissociation from dimer to monomer form, and an understanding of this process can aid in design of molecular analogs with desired properties. The practical importance of understanding insulin dimer dissociation has in turn made this system a paradigm for study of complex molecular recognition reactions. Here, we assembled a computational pipeline of methods for the study of complex molecular dynamics and applied it to understanding how the insulin dimer dissociates. This pipeline enabled us to identify collective variables that could promote dissociation with a minimum of monomeric unfolding. The resulting simulations revealed a previously unappreciated diversity of dissociation pathways with comparable free energy barriers. The limiting pathways, in which either the interfacial *α* helices separate and are solvated first (the *α* path) or the interfacial *β* strands separate and are solvated first (the *β* path), correspond to induced fit and conformational selection mechanisms when considering them from the perspective of association. The similarities in barrier heights for qualitatively different pathways makes clear the importance of achieving chemical precision in the simulations, and an error estimator that we recently introduced allowed us to do so efficiently.

Along the two limiting pathways, the elements at the dimer interface rotate relative to each other as the monomers come apart. Along the energetically preferred *α* path, the rotation allows the formation of nonnative interactions involving residues on the interfacial *α* helices; this enables the C-terminal segment of insulin B chain to detach from the nearby interfacial *α* helix, thereby coupling monomeric unfolding to unbinding. No such unfolding is observed along the *β* path. The diversity of paths that we observe encompasses paths previously observed in simulation studies;^30,33^ in this sense, our work reconciles seemingly discordant results in the literature.

Molecular simulations can guide the design and interpretation of experiments. The pathways that we observe provide a microscopic picture of a partially unfolded intermediate with conserved secondary structure previously suggested by T-jump experiments, though differences in conditions between the simulations and experiments make this picture tentative. With a view towards obtaining additional experimental constraints on the mechanism, we use the simulations to test possible sites for isotopic labeling for IR spectroscopy experiments. We predict that two sites in particular, Phe^B24^ and Glu^B13^, should enable sensitive characterization of the solvation of the interfacial *α* helices and *β* strands. Our results thus provide insight into how to pursue the next generation of experiments to achieve residue-level resolution of the dissociation mechanism.

## Methods

With a view toward providing a quantitative interpretation of experimental observations, we model insulin in solution at atomic resolution (System Setup and Equilibration), such that dimer dissociation occurs on timescales that are long compared with the molecular dynamics timestep. Consequently, both efficient sampling and informative analysis rely on identifying collective variables (CVs) that capture the slowest relaxing degrees of freedom involved in dimer dissociation. To this end, we tested many combinations of CVs for their ability to enable us to harvest reactive events (String Method and Collective Variable Selection). We found that CVs based on selected intermolecular contacts in the dimer enabled us to harvest reactive events without the addition of restraints to prevent monomer unfolding, and we improved the contact definition over the course of the study, as we gained understanding of the system (Definition of Contacts). Care was taken to converge the potential of mean force (free energy) as a function of those CVs (Adiabatic-Bias Molecular Dynamics; Replica Exchange Umbrella Sampling; Eigenvector Method for Umbrella Sampling and Adaptive Sampling). We were able to trace multiple minimum free energy paths with comparable barriers on that surface, which we validated as stable through further simulations (Finding and Confirming Energetically Favorable Paths). Finally, we computed simulated infrared spectra to guide the design of further experiments (FTIR and 2DIR Simulation). We describe each of these steps in detail below in the parenthetically indicated sections.

### System Setup and Equilibration

The system was modeled with the CHARMM36m force field.^47–49^ All simulations were performed using GROMACS 5.1.4, ^50^ and the system was prepared using CHARMM-GUI 2.1.^51,52^ Unless otherwise noted, simulations were carried out in the isochoric isothermal (NVT) ensemble at 303.15 K using a Langevin thermostat^53^ with a 2 fs timestep and a friction constant of 0.5 ps^*−*1^ applied to all atoms. All bonds to hydrogen atoms were constrained using the LINCS algorithm^54^. Periodic boundary conditions were employed and the particle-mesh Ewald method^55^ was used to calculate electrostatic forces with a cutoff distance of 1.2 nm. The Lennard-Jones interactions were smoothly switched off from 1.0 to 1.2 nm through the built-in GROMACS force-switch function. All molecular visualizations were done in VMD,^56^ and residue interaction energies were calculated using its NAMDenergy plug-in.^57^

The dimer structure was based on the human insulin crystal structure (PDB ID 3W7Y). ^58^ To fully equilibrate the system at the desired temperature and pressure, the protein was solvated, equilibrated with restraints in both the NVT and isobaric isothermal (NPT) ensembles, and then equilibrated restraint-free in the NVT ensemble. Specifically, hydrogens were added to the PDB structure, and it was solvated in a cubic box of size (8 nm)^3^ using TIP3P water^59^; 48 K^+^ and 44 Cl^*−*^ ions were added to neutralize the system and bring it to a concentration of 150 mM KCl^60^. There was a total of 48,260 atoms. The system was energetically minimized using the steepest descent method, until the maximum force felt by the system was below 1000 kJ/mol nm. The system was then equilibrated for 100 ps in the NVT ensemble with a 1 fs timestep, followed by 10 ns in the NPT ensemble at 1 bar using the Parrinello-Rahman barostat,^61^ with a 2 fs timestep and time constant of 5.0 ps. For the energy minimization and equilibration above, harmonic restraints were used to stabilize the positions of all non-hydrogen protein atoms. The system was equilibrated further for 1 ns in the NPT ensemble without position restraints, and the average box size was determined to be (7.82 nm)^3^. This box size was used for all further simulations. The system was equilibrated once more without position restraints for 1 ns in the NVT ensemble. The resulting equilibrated structure, with a root-mean-square deviation (RMSD) of 2.02 Å from the 3WY7 crystal structure, was used to initialize further simulations as described below.

### Definition of Contacts

Throughout the simulations in this work, relevant inter-residue *α* carbon distances were transformed by contact functions that smoothly vary between a small range of values. This was done to improve computational control in various methods, providing a consistent scale for biasing variables as distances varied between small and large values. The contact functions were tuned to each method, and as we learned more about the structural features of the dissociation. Specifically, we used the following three contact definitions to transform the distance between *α* carbons of residues *i* and *j* (*d*_*ij*_):

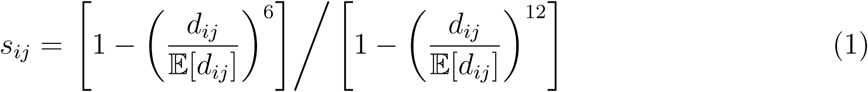

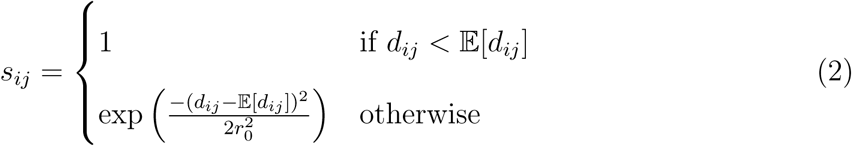

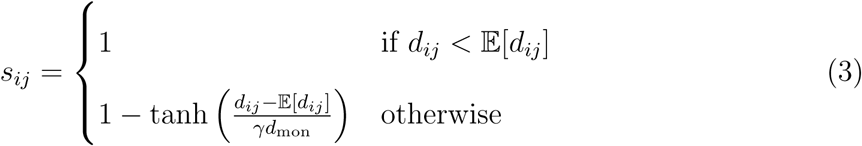

In the above equations, 𝔼 denotes an equilibrium average, and the average distance 𝔼[*d*_*ij*_] was measured for each contact pair from a 5 ns simulation, initialized from the equilibrated dimer structure. The definition in Equation 1 was used for the initial driving and string method simulations used to discover collective variables. The definition in Equation 2 was used for the umbrella sampling calculations, as it provided a gentler bias near the dimer state. In this definition, *r*_0_ is a parameter that sets the location of the inflection point of the Gaussian transformation. This parameter was set to 0.6 nm to ensure that there were at least three layers of water between all monomeric residues at dissociation, based on visual inspection. Finally, the definition in Equation 3 was used for the final string calculations used to verify the stability of our observed low-energy paths, as it provided better resolution near the dimer state. In this definition, *d*_mon_ is the average residue pair distance that corresponds to the monomeric state, again chosen so that at least three layers of water separate the residues of interest. This was thus set to be 2.2 nm for the *α* contacts and 2.0 nm for the *β* contacts (see Results). The parameter choice *γ* = 0.65 tunes the sizes of the monomeric and dimeric states in the 2D contact space.

### String Method and Collective Variable Selection

Umbrella sampling is predicated on finding a small number of collective variables (CVs) that capture the slowest relaxing degrees of freedom relevant to the process of interest. To determine reasonable CVs for insulin dimer dissociation, we tested various combinations of CVs for their ability to drive dissociation in steered molecular dynamics simulations (SMD)^62^ and then selected CVs that preserved the ability to distinguish refined dissociation paths obtained from the string method,^63,64^ discussed in further detail below. The CVs explored were based on interacting pairs of residues with high differential solvent accessible surface area (SASA) between dimer and monomer states (Supplemental Table S1, where the apostrophe differentiates residues on one monomer from residues on the other). These included the aromatic triplet in the interfacial *β* sheet, which was previously identified as important for dimer stability. ^17,19,65^

Distances between the C_*α*_ atoms of these residue pairs were computed and transformed using Equation 1 as described above. Constant velocity SMD simulations, in which harmonic restraints were used to advance random subsets of *s*_*ij*_ from 0.5 to 0.0, were used to drive the system from the equilibrated dimer structure to the dissociated state. By visual analysis, we selected dissociation paths that both led to complete dissociation and did not involve significant unfolding of the monomers since there is limited experimental evidence for extensive loss of secondary structure.^25,29^ We also only selected paths that had maximum free energies within the range of previous simulations.^19,30^ These were used to initialize string method simulations in the 22-dimensional space of intermonomeric contacts described above (Supplemental Table S1). Convergence of strings during the simulations was computed by the Hausdorff distance metric between the current string iteration and the initial string, and simulations were run until this distance metric did not change significantly.^66^

To choose a small number of CVs sufficient to describe the dissociation, we clustered the initial and final strings in the space of the first few coordinates obtained from applying the diffusion maps method^67,68^ to their images. We then sought physically interpretable CVs that preserved the clusters, which led to selection of two average contact functions (averaging performed after the transformation), one for three residue pairs at the *β* sheet interface 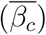 and one for seven residues at the *α* helical interface 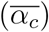. These are detailed in Supplemental Table S1 and Figure 2. These residue pairs are consistent with important interfacial interactions identified in a recent steered molecular dynamics study.^17^ We denote the average of the raw distances associated with 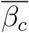 and 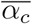, used for visualization, by 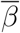 and 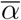, respectively.

### Adiabatic-Bias Molecular Dynamics (ABMD)

To initialize sampling, 49 independent ABMD^69^ molecular dynamics simulations (using the PLUMED 2.3 wrapper for GRO-MACS^70–72^) were used to drive the system from the dimer to a 7 × 7 grid of points evenly covering the 2D CV space of 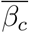 and 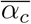. ABMD is similar to SMD but ratchets the system to its target following unbiased fluctuations along the CVs; we came to prefer it to SMD because we found that SMD but not ABMD resulted in melting of the interfacial *α* helices. That said, because ABMD relies on unbiased fluctuations, we found that the initial 49 simulations did not adequately sample the space close to the 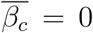 and 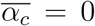 axes. We thus performed 13 extra simulations to drive the system to supplementary points near where either one or both of 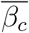 or 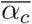 went to zero. This driving, whose bias was applied on the 10 individual distances associated with 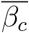 and 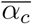, was repeated with force constants of 1000, 3000, and 5000 kJ/(mol nm), generating a database of trajectories that covered all of the relevant average contact space.

### Replica Exchange Umbrella Sampling (REUS)

The window centers for the umbrella sampling were distributed on a logarithmically spaced grid in the space of 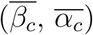 with the expectation that the initial steps of the dissociation would involve larger changes in free energy, as seen in Figure 8A. The force constants, *k*, for the harmonic biases associated with each window were described by the following equation, adapted from an expression derived by Im and coworkers:^73,74^

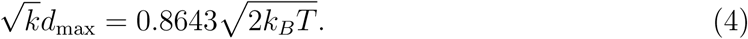

Here, as the windows are unevenly spaced, *d*_max_ refers to the maximum distance between adjacent window centers. Additional weak upper walls (half harmonic potentials with *k* = 50 kJ/(mol nm), turned on at 3 nm for each distance) were placed on each distance to prevent artificial interactions across periodic boundaries. To initialize each window, the ABMD-generated structure that was nearest to each minimum was selected and equilibrated for 100 ps using the harmonic restraint described by equation 4. A 2D replica exchange procedure, in which there were exchanges between windows, ^75^ was implemented by taking advantage of the built-in functionality of GROMACS. Each of the 784 windows were simulated for 100 ps, with exchanges attempted at 1 ps intervals only between adjacent windows in the same row. The windows were then simulated for an additional 100 ps, with exchanges attempted every 1 ps between adjacent windows in the same column. This procedure was repeated for a total of 5 ns of simulation time per window, with structures saved every 5 ps. In this way, replicas were exchanged via all nearest neighbors across the entire lattice of windows. Exchange probabilities between 10-50% were achieved depending on the specific window pairs being swapped.

### Eigenvector Method for Umbrella Sampling (EMUS) and Adaptive Sampling

The 5 ns of sampling per window was combined to generate a potential of mean force (PMF) by using the Eigenvector Method for Umbrella Sampling (EMUS).^34^ Once the PMF was created, we wanted to investigate whether the PMF was converged and, if not, add additional sampling selectively where it would be most effective. To do this, EMUS was used to estimate the asymptotic variance of replica exchange umbrella sampling simulations. However, because of the replica exchange, assumption VII.3 of ref. 34, namely that sampling in each window is independent, does not hold for our study. Nonetheless, one can still apply Lemma VII.2 in ref. 34 to derive a central limit theorem for EMUS with replica exchange by casting sampling over all windows as a Markov chain; in this case, the asymptotic covariance matrix, Σ, is not block diagonal. This leads to a definition of the asymptotic variance for arbitrary averages that one can approximate by an expectation of integrated autocovariances over the sampled data. For details, see the Supplemental Information.

The contact space PMF is shown in Figure 8B, with its associated asymptotic variance in Figure 8C. The area of highest asymptotic variance in the PMF was identified, and is marked by a red box in Figure 8C. Using the process described in ref. 34, the per-window error contributions to this region were determined; although these include only the error we would observe if the off-diagonal blocks of Σ were zero, we believe they are sufficient to diagnose the behavior of the umbrella sampling scheme. These contributions are shown in Figure 8D, and reveal a J-shaped region of windows which contribute the most to the asymptotic variance of the region marked in Figure 8C. These windows were then identified as areas to add additional sampling. This additional sampling used a similar procedure as described above for the initial replica exchange simulations, with the sampling and proposed exchanges restricted to the bottom-most five rows and right-most 5 columns in Figure 8D. At 1 ns intervals, this additional sampling was independently processed with EMUS as follows. These data were used to compute a new PMF and its associated asymptotic variance per bin, 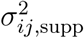. We then combined 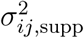 with the initial asymptotic variance, 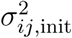, weighted by the squared ratio of simulation lengths between supplemental and total sampling, 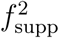:

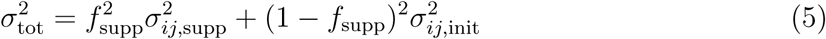

Equation 5 assumes the initial 5 ns and supplemental 5 ns of sampling are independent, which given that the autocorrelation time of the quantities needed for EMUS was 6 ps on average across the windows, is a reasonable assumption.

As seen in Figure 8E, the peak variance decreased from 0.25 to 0.12 (kcal/mol)^2^ upon the addition of 5 ns of additional sampling per window in the outlined J-shaped region. This region would not obviously be chosen in the absence of a quantitative procedure, though it can be rationalized in hindsight as corresponding to a major dissociation pathway that we characterize in detail in Results and Discussion. This illustrates how EMUS allows users to monitor the convergence of US simulations as sampling proceeds, and to adaptively identify regions of state space that are most in need of additional sampling, despite the neglect of correlations discussed above. Furthermore, EMUS allows for the calculation of PMFs in arbitrary CV spaces, not just the space in which the biasing was done, without the need for additional sampling. Examples of these PMFs are seen in Results and Discussion.

### Finding and Confirming Energetically Favorable Paths

Our analysis of dissociation is based on minimum free energy paths. Initially, seven such paths were drawn on the 2D PMF by using the lfep search algorithm.^76^ To ensure that these were stable in the 10-dimensional space of all of the contacts associated with the averages 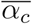 and 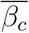, another iteration of the string method was run in this space, this time initialized from structures drawn from the REUS database along these 2D minimum free energy paths. The specific contact definition used is in Equation 3, and the strings were run until converged as measured by the Hausdorff distance, as discussed previously. The starting and ending positions of these strings are shown in Supplemental Figure S1. In particular, the *α* path shows almost no variation, and the *β* path shifts only minimally, and this shift does not change any of the molecular trends discussed in the Results. The others paths also exhibit minimal variation. This provides evidence that most of the pathways we identify and the limiting pathways in particular are indeed stable in a broader space.

### FTIR and 2DIR Simulation

Simulated IR spectra were calculated from the Fourier transform of a vibrational transition dipole time correlation function using a mixed quantum-classical model described in refs. 43 and 77. Briefly, electrostatic collective variables can be used to translate molecular-dynamics sampling of protein structure into (i) a time-dependent Hamiltonian and (ii) a transition dipole moment that describes the amide I vibrations of protein backbones; these quantities in turn can be used to calculate the dipole time correlation functions that are needed to create simulated FTIR and 2D IR spectra. Furthermore, these spectra can be calculated for both native proteins and isotope-labeled proteins.^46^ Here, we aimed to generate FTIR spectra for 50 points equally spaced along both the *α* and the *β* paths for a variety of isotope-labeled insulins. 2DIR spectra were then calculated for specific states for two isotope-labeled insulins. These spectra were then used to propose possible experiments to validate our results.

First, each path was divided into a series of 50 points, referred to as image centers. By comparing these image centers to the REUS database, 20 structures were selected to be associated with each image center. These structures were randomly drawn from the sampling that was within both 0.4 nm of each image center and the Voronoi tessellation associated with each image center. Each of these structures served as the starting point for an additional short molecular dynamics simulation consisting of 100 ps of equilibration followed by 100 ps of sampling every 20 fs. These simulations were then used to generate both the FTIR and 2DIR spectra by using the procedure outlined below.

As IR spectra correspond to manifestly quantum-mechanical vibrational transitions, our classical molecular dynamics trajectories had to be translated into a time-dependent Hamiltonian and transition dipole trajectories. To this end, we associated each amide I vibration with a site, defined by the atomic positions of the backbone amide groups (C, O, N, and H atoms). The frequency of each site was calculated using an empirical electrostatic frequency map optimized against experimental spectra of isotope-edited NuG2b protein, which evaluates the electrostatic potential value at the C, O, N, and H positions. ^78^ This potential-based map (4PN-150) has an estimated frequency uncertainty of 2.25 cm^*−*1^. When applying the map, we used modified glycine charges described previously.^78^ Additionally, we considered coupling between sites, including both through-bond mechanical coupling and through-space electrostatic coupling. Through-bond coupling between adjacent sites was generated using a density functional theory (DFT)-based nearest-neighbor coupling map, while through-space coupling was computed by a transition charge coupling map. ^79^ We did not account for vibrations from protein side-chain and terminal groups. The transition dipole of each site was assigned using the zero-field values from a DFT-based electrostatic map.^80^ When generating simulated 2DIR spectra, there are signal contributions from excited state absorption (ESA), which correspond to vibrational transitions between states with one quantum of excitation energy and those with two quanta of excitation energy. To deal with this, the corresponding two-quantum Hamiltonian and transition dipole moments were constructed using a weak anharmonic model.^43,81^

The time-dependent Hamiltonian and transition dipole trajectories were converted to simulated FTIR spectra and 2D IR spectra using a dynamic wavefunction-propagation scheme with a Trotter expansion to reduce computation time.^82,83^ The window time for calculating dipole time correlation functions was set to 2.5 ps. The anharmonicity of the amide I oscillator was set to 16 cm^*−*1^.^81^ The amide I vibrational lifetime was modeled by an ad hoc single exponential decay, with a time constant of 1.0 ps determined by transient absorption experiments of Ala–Ala. ^84^ The isotope frequency shift introduced by a ^13^C^18^O label was set to 65 cm^*−*1^.^43^ The spectra for the structures that were selected from the REUS database were uniformly averaged within each of the 50 images across both paths. This created a simulated spectrum representative of the location of each image in CV space. The isotope labeled FTIR spectra along each path, with the corresponding simulated unlabeled spectra subtracted to create difference spectra, are shown in Supplemental Figure S11. Based on these results, the data were regrouped and reaveraged as described in the Supplemental Information to create the 2DIR spectra shown in Figure 7.

## Supporting information

Representative Structures in PDB Format

## Acknowledgments

The authors thank John Strahan, Chatipat Lorpaiboon, Biman Bagchi, and Michael Weiss for helpful discussions. We thank Albert Pan, Bryan Jackson, and coworkers at D.E. Shaw for providing their trajectories for comparison. This work was supported by National Institutes of Health awards R01GM109455-01 and 5R01GM118774-02. Computations were performed on resources provided by the University of Chicago Research Computing Center, and the Extreme Science and Engineering Discovery Environment^85^ (NSF Grant ACI-1548562) Bridges (PSC) computing nodes through allocation TG-MCB180007.

## Supporting Information Available

Supplementary figures (as referenced in the text) and accompanying commentary are available, as well as coordinate files for the structures shown in Figure 3 and used throughout this work.

