## Supporting Text, Tables, and Figures for "Insulin dissociates by diverse mechanisms of coupled unfolding and unbinding"

### Supporting Information for Insulin dissociates by diverse mechanisms of coupled folding and binding

**EMUS Asymptotic Error With Replica Exchange.** To derive the expression for asymptotic variances of averages given REUS data, we follow the following procedure. The notation corresponds to that in ref. 1.

**Assumption 0.1.** *We assume:*

1. *Sampling over all windows is performed by a single Markov chain $\Xi$ whose state space is $L$ copies of the molecular phase space.*

2. *A central limit theorem holds for the convergence of each sample average $\bar{v}$ estimated in EMUS to its true value $v$, with asymptotic covariance matrix $\Sigma$. The entries in this matrix are given by*

$$\Sigma_{ij} = \frac{1}{2} \int_{-\infty}^{\infty} \mathbb{E}\left[v_i(X_t) v_j(X_0)\right] dt. \tag{1}$$

3. *The biasing functions $\psi_i$ are chosen so that $F$ is irreducible.*

Note that if we sort the averages $v$ by the window in which the average was calculated, then unlike in ref. 1, the matrix $\Sigma$ is not block diagonal. Rather it has nonzero off-diagonal blocks:

$$\Sigma = \left[\begin{array}{c|c|c} \Sigma_{11} & \Sigma_{12} & \cdots \\ \hline \Sigma_{21} & \Sigma_{22} & \cdots \\ \hline \vdots & \vdots & \ddots \end{array}\right].$$

Under these assumptions, we can still apply Lemma VII.2 in ref. 1 to derive a central limit theorem for EMUS with replica exchange.

**Theorem 0.2.** *As in ref. 1, let $B$ be a quantity of interest and $dB/d\bar{v}$ be the partial derivative of $B$ with respect to each sampled average. Under the assumptions above,*

$$\sqrt{N}\left(B\left(\bar{v}\right) - B\left(v\right)\right) \xrightarrow{\mathrm{d}} N(0, \sigma^2), \tag{2}$$

*where*

$$\sigma^2 = \frac{\partial B}{\partial \bar{v}}^T \Sigma \frac{\partial B}{\partial \bar{v}}. \tag{3}$$

To estimate $\sigma^2$ using sampled data, we write

$$\sigma^2 = \sum_{i,j} \frac{\partial B}{\partial \bar{v}_i} \Sigma_{ij} \frac{\partial B}{\partial \bar{v}_j} \tag{4}$$

$$= \sum_{i,j} \frac{\partial B}{\partial \bar{v}_i} \int_{-\infty}^{\infty} \mathbb{E}\left[v_i(X_t) v_j(X_0)\right] dt \frac{\partial B}{\partial \bar{v}_j}$$

$$= \int_{-\infty}^{\infty} \mathbb{E}\left[\left(\sum_i \frac{\partial B}{\partial \bar{v}_i} v_i(X_t)\right)\left(\sum_j \frac{\partial B}{\partial \bar{v}_j} v_j(X_0)\right)\right] dt \tag{5}$$

This is the the integrated covariance of the trajectory $\sum_i \frac{\partial B}{\partial \bar{v}_i} v_i(X_t)$.

**C-terminal detachment is correlated with solvation of the A-chain N-terminal segment and formation of nonnative contacts.** The detachment of the C-terminal segment of the B chain (Figure S9), discussed extensively in the main text, is also correlated with the increase in SASA of $\mathrm{Gly^{A1}}$-$\mathrm{Val^{A3}}$ (Figure S10). $\mathrm{Gly^{A1}}$-$\mathrm{Val^{A3}}$ are residues that are in contact with $\mathrm{Pro^{B28}}$-$\mathrm{Ala^{B30}}$ in the dimer, and these contacts are sacrificed as detachment occurs along the $\alpha$ path. This is consistent with the known binding behavior of the insulin monomer to the insulin receptor, as these $\mathrm{Gly^{A1}}$-$\mathrm{Val^{A3}}$ residues are thought to form part of the binding interface[2,3] and are not in contact with $\mathrm{Pro^{B28}}$-$\mathrm{Ala^{B30}}$ when bound to the receptor.[4,5]

Beyond the loss of contacts between $\mathrm{Pro^{B28}}$-$\mathrm{Ala^{B30}}$ and $\mathrm{Gly^{A1}}$-$\mathrm{Val^{A3}}$ along the $\alpha$ path, one can also characterize the number of contacts that the B-chain C-terminal segment makes with other residues. A subset of these are shown in Figure S4. Namely, from the left panels, one sees that along the $\alpha$ path (black), the native contacts of $\mathrm{Pro^{B28}}$-$\mathrm{Ala^{B30}}$ with $\mathrm{Gly^{B'20}}$-$\mathrm{Gly^{B'23}}$ and $\mathrm{Gly^{A1}}$-$\mathrm{Val^{A3}}$ are lost. The rotation of the interface associated with both the $\alpha$ and $\beta$ paths moves the B-chain C-terminal segment away from these both $\mathrm{Gly^{B'20}}$-$\mathrm{Gly^{B'23}}$ and $\mathrm{Gly^{A1}}$-$\mathrm{Val^{A3}}$.

The right plots, on the other hand, show that some specific nonnative interactions are formed along the $\alpha$ path, but not along the $\beta$ path. Along the $\alpha$ path, the B-chain C-terminal segment starts to interact with the Tyr$^{B'16}$ on the $\alpha$ helix of the other monomer when $\overline{\alpha} \approx 1.15$ nm. In the same region, Ser$^{B9}$ and Ser$^{B'9}$, serines on opposite interfacial helices, start to interact and form a nonnative contact. Interestingly, comparing this to the core SASA shown in the main text Figure 5A, one sees that the core SASA increases sharply after the Ser$^{B9}$ residues lose contact with one another, around $\overline{\alpha} \approx 1.35$ nm. This is also the area where significant detachment starts to occur along the $\alpha$ path. It is possible that the combination of the interfacial rotations and detachment exposes the hydrophobic core of the dimer.

**Relation to available insulin therapeutics.** To replicate the biphasic activity of insulin *in vivo*, both fast-acting and slow-acting insulin therapeutics have been developed. Two of the most common fast acting therapeutics, lispro (Pro$^{B28}$Lys$^{B29}$ $\rightarrow$ Lys$^{B28}$Pro$^{B29}$, Humalog[6]) and aspart (Pro$^{B28}$ $\rightarrow$ Asp$^{B28}$, NovoLog[7]) involve mutation of C-terminal residues of the B chain to destabilize the dimer and favor the monomer. This presumably allows for more rapid and reliable uptake of glucose after injection, as the monomer is the species that binds to the receptor. These mutations have been hypothesized to reduce the interaction of Pro$^{B28}$-Ala$^{B30}$ of one monomer with the $\beta$ turn of the other monomer (Gly$^{B'20}$-Gly$^{B'23}$), either through reduced van der Waals attractive forces (lispro and aspart) or the addition of a repulsive electrostatic interaction (aspart).[8] To the best of our knowledge, no computational work has yet been done to explore these hypotheses as they relate to the mechanism of dissociation. In the previous section, it was described how, along both pathways, the native contacts between Pro$^{B28}$-Ala$^{B30}$ and both Gly$^{B'20}$-Gly$^{B'23}$ and Gly$^{A1}$-Val$^{A3}$ are broken. It has been suggested[6,8] that one or both of the therapeutic mutations could reduce the energetic penalty for breaking these native contacts, specifically the contacts with Gly$^{B'20}$-Gly$^{B'23}$. To explicitly investigate this in our simulations of wildtype insulin, the sum of interaction energies of Pro$^{B28}$-Ala$^{B30}$ and both Gly$^{B'20}$-Gly$^{B'23}$ and Gly$^{A1}$-Val$^{A3}$ was computed (Figure

S5).

**$\beta$ turn angle and disorder of the $\beta$ turn also varies between dissociation path.** We defined the $\beta$ turn angle, $\Phi_t$, as the angle between the geometrical centers of the backbone atoms in $Glu^{B13}$ , $Gly^{B20}$, and $Phe^{B24}$ to characterize the $\beta$ turn. A larger $\beta$ turn angle corresponds to a wider turn. We also calculated the RMSD of the $\beta$ turn from its conformation in the dimer by fitting the conserved $\alpha$ helix of the B chain to each monomeric unit, then measuring the RMSD to the solvated crystal structure. The average of both the $\beta$ turn angle and the RMSD are shown in Figure S8.

The $\beta$ turn angle increases initially and is much wider along the $\alpha$ path than along the $\beta$ path. The RMSD also increases as the $\alpha$ helices separate. This increase in RMSD happens before the $\beta$ sheets are broken for the $\alpha$ path, and after the $\beta$ sheets are broken for the $\beta$ path. The different trends in $\beta$ turn angle and RMSD between paths suggest that these residues on the $\beta$ turn might be useful candidates for isotopic labeling for 2DIR experiments aimed at distinguishing dissociation pathways.

**FTIR to identify labels and solvated species.** Here we describe our simulations of FTIR spectra to map the effects of isotopic labels with considerably less computational cost than simulations of 2DIR spectra. As noted in the main text, we computed 50 such spectra along each of the two limiting paths, and each was converted into a difference spectrum by subtracting the unlabeled insulin spectrum at that point along the path from the labeled spectrum. The intensities of these difference spectra were converted into a colormap, with orange being a positive change, and blue being a negative change. This allows one to see the peaks associated specifically with the isotope label. These were then stacked and viewed as a function of path progress along both the $\alpha$ and $\beta$ paths. Thus, when a red feature shifts to a lower frequency, this corresponds to a redshift in the simulated labeled spectra. These series of spectra, for the $Phe^{B24}$ and $Glu^{B13}$ isotope-labeled insulins, are shown in Figure S11. We simulated all possible constructs with a single interfacial residue isotopically labeled (between $Ser^{B9}$ and $Ala^{B30}$). The labels on our identified $\alpha$ and $\beta$ contacts showed the most significant redshifts; of these $Phe^{B24}$ and $Glu^{B13}$ exemplified these effects.

From these spectra, we identified the ranges of path progress where the redshift (and correlated loss in intensity) first occurs, presumably due to either solvation or change in local secondary structure. These areas are marked areas in Figure S11. The left panels show the SASA of the backbone carbonyl group associated with the isotopically labeled residue (either $Phe^{B24}$ or $Glu^{B13}$ ). One sees that these marked regions begin once the SASAs for the corresponding backbone carbonyl groups increase from the dimer (with SASAs near 0) to 60% of the monomeric average. One can thus conclude that the redshifting and peak broadening in these FTIR spectra correlates with the solvation of the backbone.

Table S1: Types and descriptions of contacts with high differential SASA between dimer and monomer, with primes indicating intermonomer interactions.

| Class | Residue Pairs | Description |
|---|---|---|
| $\beta$ Contacts | $\mathrm{Phe}^{\mathrm{B}24}$:$\mathrm{Tyr}^{\mathrm{B}'26}$, $\mathrm{Phe}^{\mathrm{B}25}$:$\mathrm{Phe}^{\mathrm{B}'25}$, $\mathrm{Tyr}^{\mathrm{B}26}$:$\mathrm{Phe}^{\mathrm{B}'24}$ | $\mathrm{Phe}^{\mathrm{B}24}$ and $\mathrm{Tyr}^{\mathrm{B}26}$ form hydrogen bonds at the $\beta$ sheet interface side chains make up part of the hydrophobic core of the dimer. $\mathrm{Phe}^{\mathrm{B}25}$ is also part of the $\beta$ sheet, but its side chain is not part of the hydrophobic core. |
| $\alpha$ Contacts | $\mathrm{Ser}^{\mathrm{B}9}$:$\mathrm{Glu}^{\mathrm{B}'13}$, $\mathrm{Ser}^{\mathrm{B}9}$:$\mathrm{Tyr}^{\mathrm{B}'16}$, $\mathrm{Val}^{\mathrm{B}12}$:$\mathrm{Tyr}^{\mathrm{B}'16}$, $\mathrm{Glu}^{\mathrm{B}13}$:$\mathrm{Ser}^{\mathrm{B}'9}$, $\mathrm{Glu}^{\mathrm{B}13}$:$\mathrm{Glu}^{\mathrm{B}'13}$, $\mathrm{Tyr}^{\mathrm{B}16}$:$\mathrm{Ser}^{\mathrm{B}'9}$, $\mathrm{Tyr}^{\mathrm{B}16}$:$\mathrm{Val}^{\mathrm{B}'12}$ | $\mathrm{Ser}^{\mathrm{B}9}$, $\mathrm{Val}^{\mathrm{B}12}$, $\mathrm{Glu}^{\mathrm{B}13}$ , $\mathrm{Tyr}^{\mathrm{B}16}$ are residues from the $\alpha$-helical portion of the B chain that are at least partially buried in the dimer. $\mathrm{Val}^{\mathrm{B}12}$ and $\mathrm{Tyr}^{\mathrm{B}16}$ are part of the hydrophobic core. |
| Cross Contacts | $\mathrm{Val}^{\mathrm{B}12}$:$\mathrm{Phe}^{\mathrm{B}'24}$, $\mathrm{Tyr}^{\mathrm{B}16}$:$\mathrm{Tyr}^{\mathrm{B}'26}$, $\mathrm{Phe}^{\mathrm{B}24}$:$\mathrm{Val}^{\mathrm{B}'12}$, $\mathrm{Tyr}^{\mathrm{B}26}$:$\mathrm{Tyr}^{\mathrm{B}'16}$ | Buried residues across the dimer interface that pair $\alpha$ helices with $\beta$ strands and vice versa. |
| Intermittent Contacts | $\mathrm{Val}^{\mathrm{B}12}$:$\mathrm{Glu}^{\mathrm{B}'13}$, $\mathrm{Glu}^{\mathrm{B}13}$:$\mathrm{Val}^{\mathrm{B}'12}$, $\mathrm{Glu}^{\mathrm{B}21}$:$\mathrm{Pro}^{\mathrm{B}'28}$, $\mathrm{Gly}^{\mathrm{B}23}$:$\mathrm{Thr}^{\mathrm{B}'27}$, $\mathrm{Thr}^{\mathrm{B}27}$:$\mathrm{Gly}^{\mathrm{B}'23}$, $\mathrm{Pro}^{\mathrm{B}28}$:$\mathrm{Glu}^{\mathrm{B}'21}$ | $\mathrm{Thr}^{\mathrm{B}27}$ and $\mathrm{Pro}^{\mathrm{B}28}$ are relatively disordered residues adjacent to the $\beta$-sheet that form intermittent contacts with $\beta$ turn residues $\mathrm{Glu}^{\mathrm{B}21}$ and $\mathrm{Gly}^{\mathrm{B}23}$. $\mathrm{Val}^{\mathrm{B}12}$ and $\mathrm{Glu}^{\mathrm{B}13}$ are on the interfacial $\alpha$ helices and also only interact intermittently. |
| Internal Contacts | $\mathrm{Val}^{\mathrm{B}12}$:$\mathrm{Tyr}^{\mathrm{B}26}$, $\mathrm{Glu}^{\mathrm{B}13}$:$\mathrm{Phe}^{\mathrm{B}24}$ | Internal contacts within a monomer. |

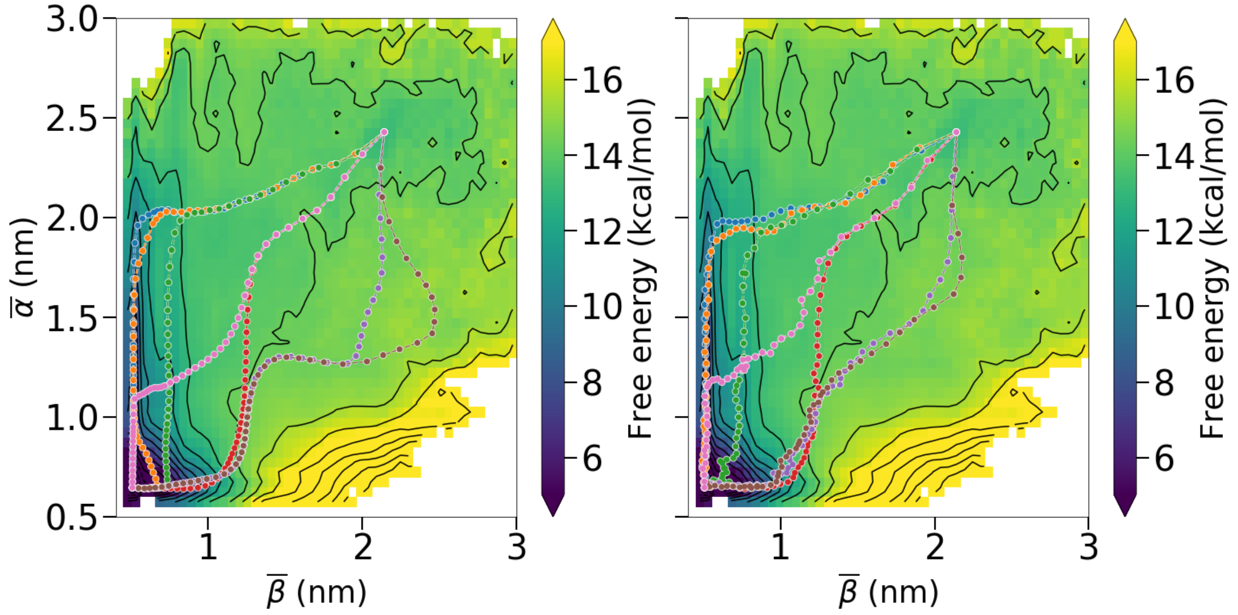


Figure S1: Using the string method to confirm stability of dissociation paths. (Left) Minimum free energy paths identified by using the LFEP algorithm from ref. 9. All the paths shown have a maximum free energetic barrier within 4 $k_BT$ of the others. These paths were used as initializations for the string method. (Right) The converged strings after further refinement with the string method. Comparing with the left panel, the orange path has collapsed to the $\alpha$ path, and the purple and brown paths have shifted slightly from their initial positions. Note that while the purple path, which represents the $\beta$ path, has shifted slightly in CV space, this does not affect the molecular trends discussed in the main text. Overall, the stability of these paths provides evidence that the averaging reducing the 10 interfacial distance dimensions to $\overline{\beta}$ and $\overline{\alpha}$ does not sacrifice mechanistic information.

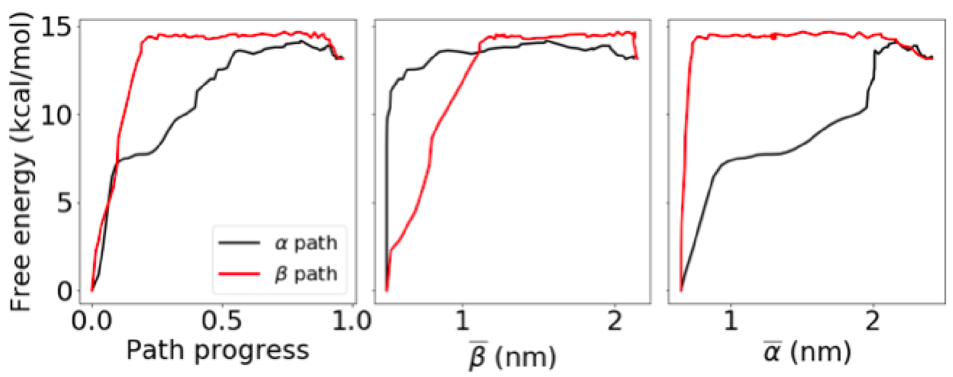


Figure S2: 1D cuts of our $\alpha$ and $\beta$ paths as functions of path progress (left), $\overline{\beta}$ (middle), and $\overline{\alpha}$ (right).

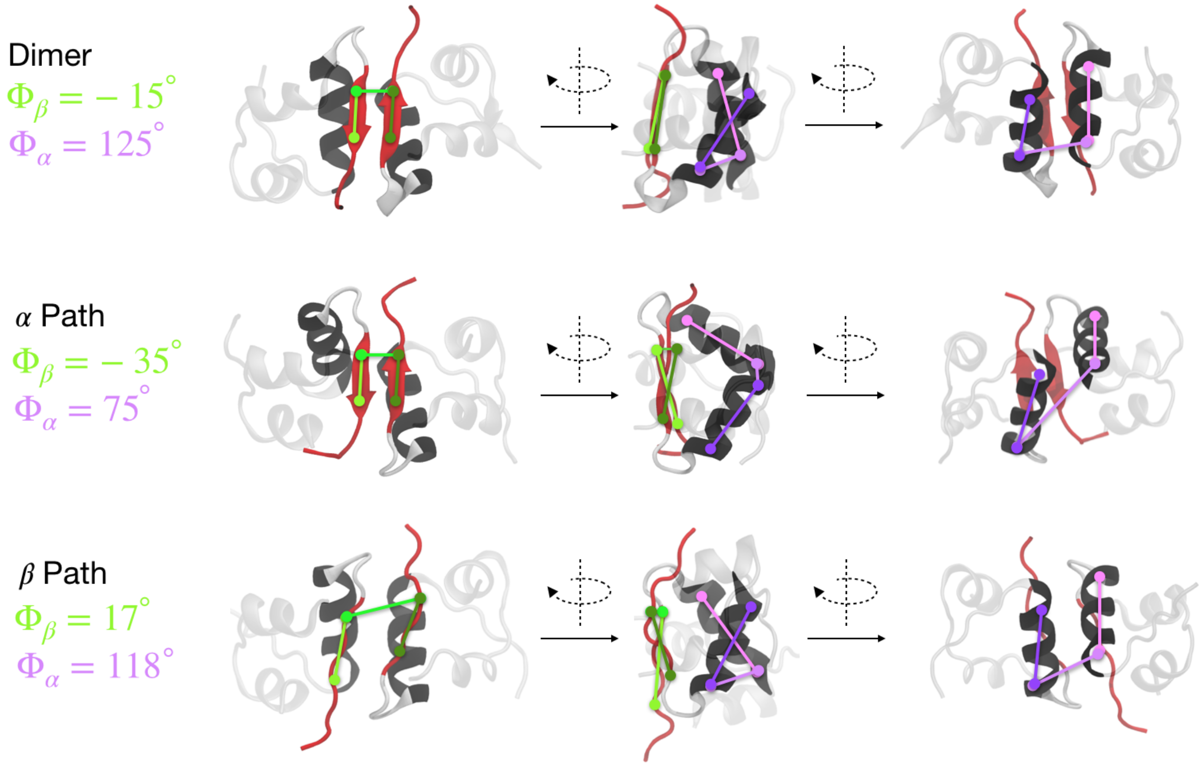


Figure S3: Structures representing the dimer (top), initial steps along the $\alpha$ path (middle), and initial steps along the $\beta$ path (bottom), with lines superimposed to show the interfacial pseudodihedral angles, $\Phi_\beta$ (green) and $\Phi_\alpha$ (purple). For these lines, a darker color in the side projection means the residues are in front while a lighter color means they are behind.

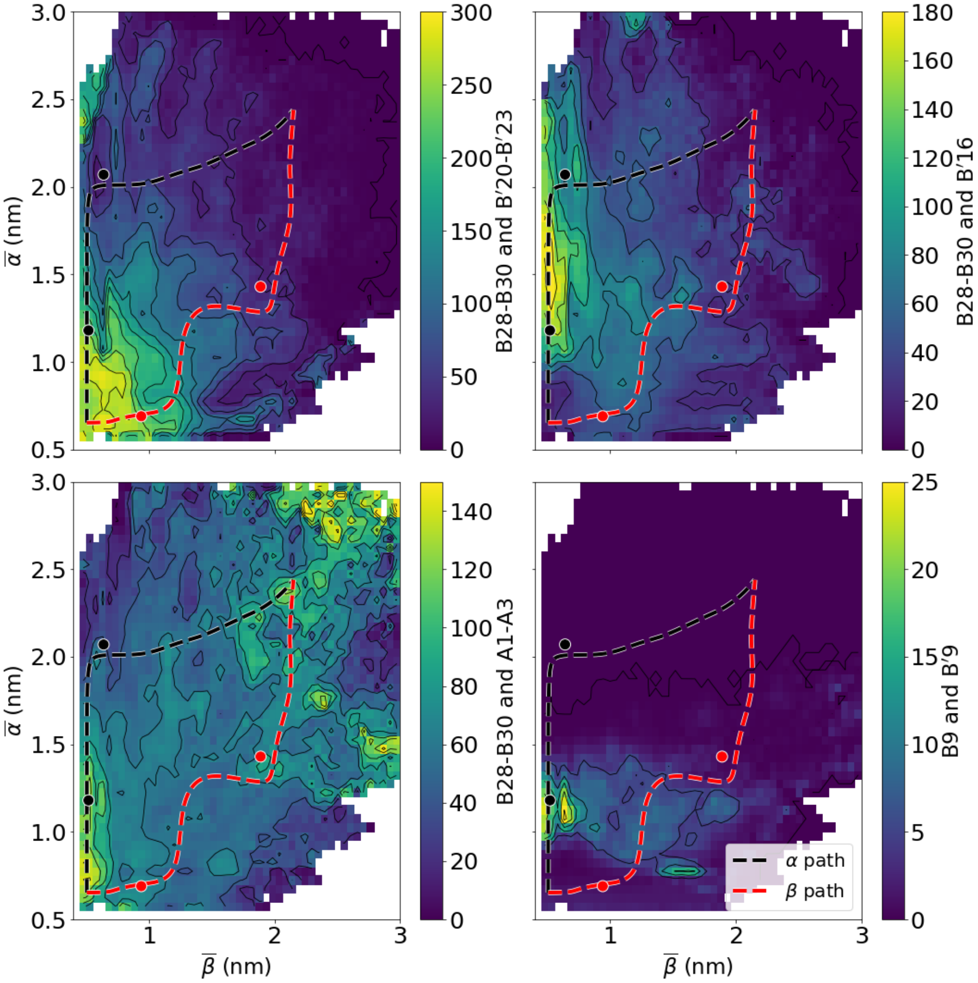


Figure S4: Averages of the number of native (left) and nonnative (right) contact pairs, with the specific pair given by the scale bar labels, as a function of $\overline{\alpha}$ and $\overline{\beta}$. The left plots show that along the $\alpha$ path, contacts between Pro$^{B28}$-Ala$^{B30}$ are lost with both Gly$^{B'20}$-Gly$^{B'23}$ (top) and Gly$^{A1}$-Val$^{A3}$ (bottom). The right plots show that along the $\alpha$ path, nonnative contacts start to form between Pro$^{B28}$-Ala$^{B30}$ and Tyr$^{B'16}$ (top), and between Ser$^{B9}$ and Ser$^{B'9}$.

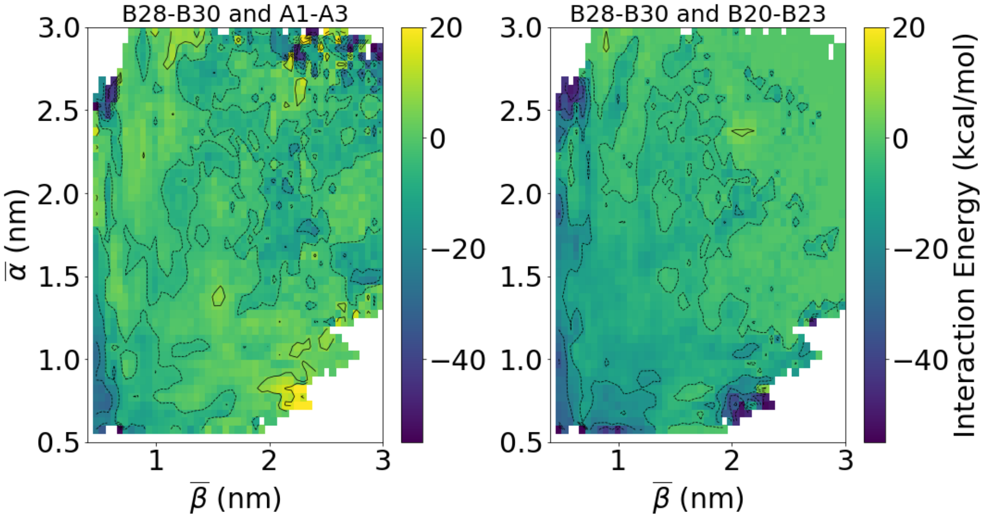


Figure S5: Average total interaction energies of Pro$^{B28}$-Ala$^{B30}$ with Gly$^{A1}$-Val$^{A3}$ (left) and Gly$^{B'20}$-Gly$^{B'23}$ (right). Contour lines shown every 10 kcal/mol. Both of these interactions stabilize the dimer state (lower left corner of both panels).

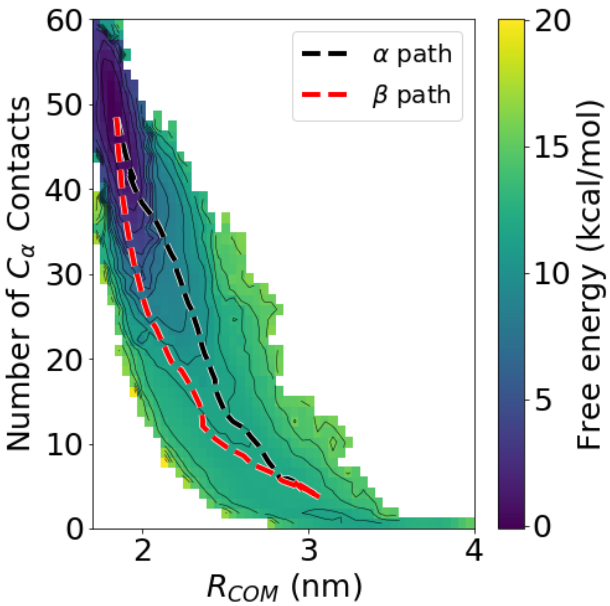


Figure S6: PMF as a function of the center of mass distance between the two monomers ($R_{COM}$) and number of interfacial C$_\alpha$ contacts (cutoff 7 Å,) the coordinates used by Bagchi and coworkers in refs. 10 and 11.

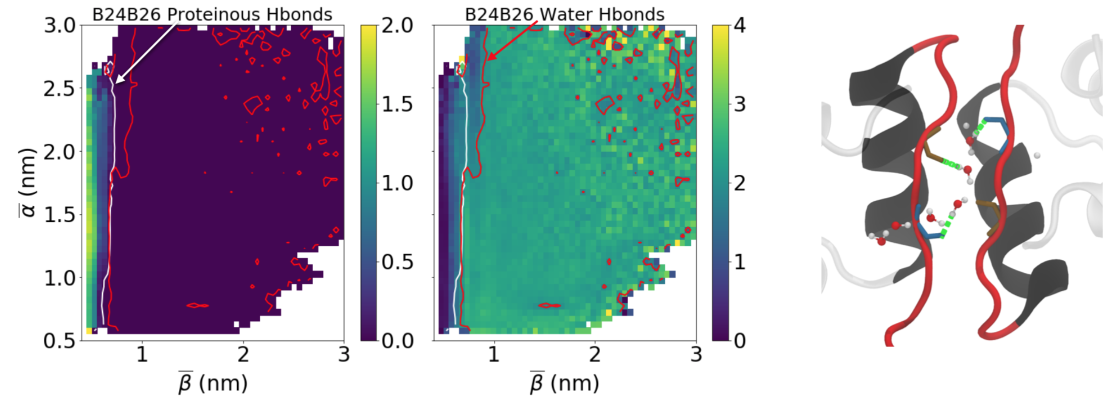


Figure S7: Characterizing interfacial hydrogen bonding. (Left) The average number of protein-protein hydrogen bonds on the beta sheet interface averaged on the PMF, compared to (middle) the average number of hydrogen bonds between those same residues and water. The white contour represents when the protein-protein hydrogen bond character drops to 2% of its initial value, while the red contour shows the point where on average 2 hydrogen bonds have been formed with water on the interface. These contours correlate well in CV space. On the right is a representative structure showing this solvation, with hydrogen bonds between the interfacial residues and water shown in green.

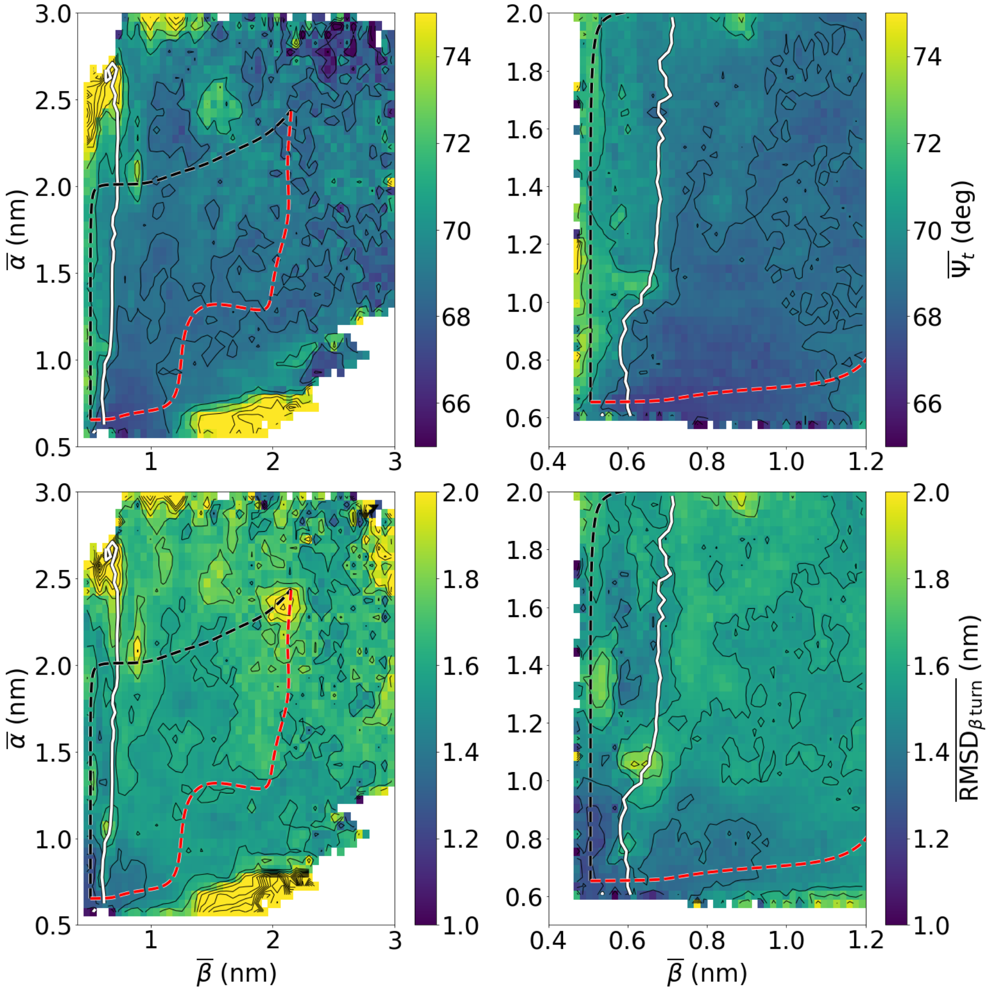


Figure S8: Plots showing the behavior of the interfacial $\beta$ turn during the dissociation. The average $\beta$ turn angle (top) and $\beta$ turn RMSD (bottom) as a function of $\overline{\alpha}$ and $\overline{\beta}$, and zoomed in to the near-dimer regime on the right. On all graphs, the $\alpha$ and $\beta$ paths are superimposed, as well as the white contour that signifies the solvation of the $\beta$ interface. Here, we see the $\beta$ turn angle increasing along the $\alpha$ path but not along the $\beta$ path. Also, the $\beta$ turn RMSD increases before the $\beta$ sheets are broken along the $\alpha$ path, while this only occurs after the $\beta$ sheets are broken for the $\beta$ path.

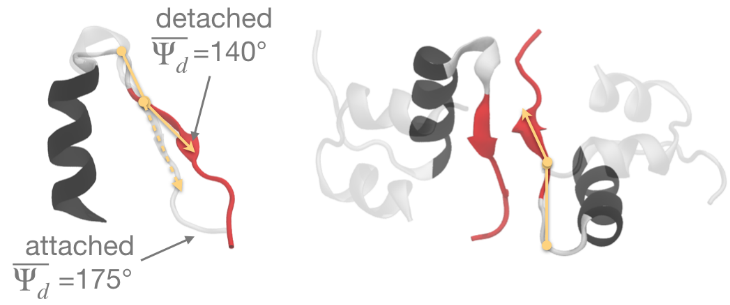


Figure S9: Structures showing detachment. (Left) The detached and attached species overlaid, with the detachment angle explicitly overlaid on top of the structure. (Right) This same detached structure, but now showing the entire dimeric species.

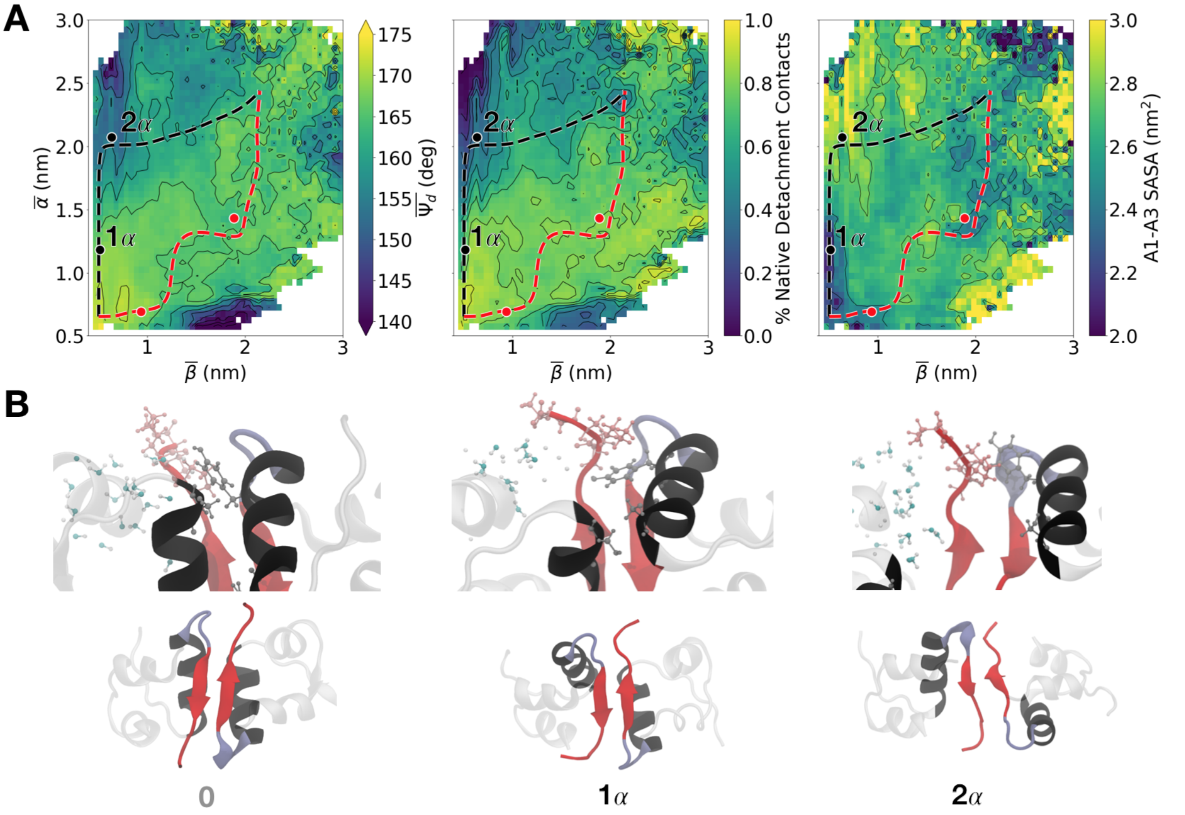


Figure S10: Detachment is correlated with solvation of the N-terminal segment of the A chain. (A) Averages of the detachment angle $\Psi_d$(left), percentage of native contacts between Pro$^{B28}$-Ala$^{B30}$ and the nearby $\alpha$ helix of the same monomer (middle), and the SASA for Gly$^{A1}$-Val$^{A3}$ of the same monomer. The similarity between the left and middle plots suggests that the detachment angle is an effective measure of the C-terminal segment moving away from the $\alpha$ helix it is normally tucked against in the native monomeric unit. Furthermore, the similarity to the rightmost plot shows that the solvation of Gly$^{A1}$-Val$^{A3}$ is correlated to the large detachment of the B chain 's C-terminal segment in the same monomeric unit. (B) Structures showing how the detachment is coupled to the solvation of Gly$^{A1}$-Val$^{A3}$.

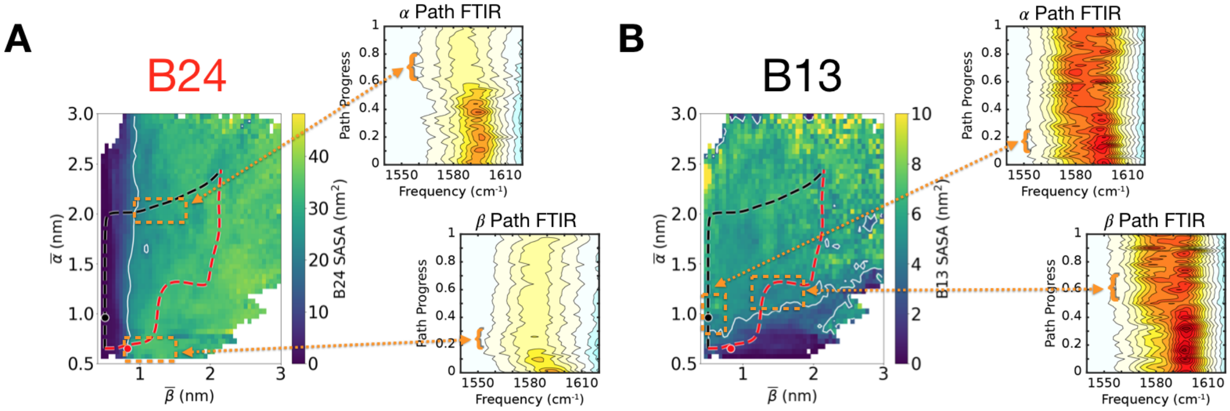


Figure S11: Relation between backbone solvation and simulated FTIR spectra. (A) The backbone carbonyl SASA for Phe$^{B24}$, with the white contour showing where the SASA increases to 60% of the monomeric average (contour at 19.7 nm$^2$, monomeric average at 32.8 nm$^2$). The two Phe$^{B24}$-labeled FTIR plots correspond to simulations along the $\alpha$ (top) and $\beta$ paths (bottom), with the y-axis being the path progress, or the fractional distance along each specific path. Each value of y corresponds to one FTIR simulation - a FTIR spectra was generated by combining 20 simulations started from a point at that specific value of path progress, and a difference was taken between that isotope-labeled simulated spectrum and the corresponding unlabeled simulated spectrum. 50 such difference spectra were created, and stacked such that the color represents the intensity of the difference. For each path, the spectra that first demonstrated the expected redshift were identified, and then those areas of path space were selected as the solvated species for future study (orange boxes on the SASA graph). (B) Similar graphs for the Glu$^{B13}$ -labeled insulin, showing the backbone carbonyl SASA for Glu$^{B13}$ . The contour is again shown where the SASA increases to 60% of the monomeric average (contour at 4.1 nm$^2$, monomeric average of 6.8 nm$^2$).
