## Supplementary material for "Insulin dissociates by diverse mechanisms of coupled unfolding and unbinding": Representative Structures in PDB Format: README

The files in this directory are the coordinate files for the structures shown
in Figure 3 of "Insulin dissociates through diverse mechanisms of coupled 
folding and binding". The entire system (protein, solvent, and ions) are
included. 

Regarding the naming convention, the names of the files match the labels in Figure 3, with 'alpha' substituted with 'a' and 'beta' substituted for 'b'. 

If you have any questions, contact Aaron Dinner at. 
3/12/2020 
